# Condensin I folds the *C. elegans* genome

**DOI:** 10.1101/2022.06.14.495661

**Authors:** Moushumi Das, Jennifer I. Semple, Anja Haemmerli, Valeriia Volodkina, Janik Scotton, Todor Gitchev, Ahrmad Annan, Julie Campos, Cyril Statzer, Alexander Dakhovnik, Collin Y. Ewald, Julien Mozziconacci, Peter Meister

## Abstract

The Structural Maintenance of Chromosomes (SMC) complexes, cohesin and condensins, are named for their roles in separating and compacting chromosomes during meiosis and mitosis. Recent data from mammalian cells have revealed additional functions for cohesin, including folding the interphase genome into loops and domains. However, it remains unclear what determines genome folding in holocentric species. To address this question, we systematically and acutely inactivated each SMC complex. Surprisingly, we found that, in contrast to mammals, condensin I is the major long-range genome loop extruder, while cohesin only creates small loops. Specifically, loss of condensin I led to genome-wide decompaction, chromosome mixing, and the disappearance of topologically associating domain (TAD) structures, while reinforcing fine-scale epigenomic compartments. Strikingly, inactivating condensin I and its X-specific variant condensin I^DC^ from the X chromosomes revealed the existence of a third compartment that groups together a subset of previously characterized loading sites for condensin I^DC^ and binding sites for the X-targeting complex SDC. Although the inactivation of cohesin, condensin II, and condensin I/I^DC^ led to minor transcriptional changes for all autosomes, removing condensin I/I^DC^ from the X chromosome resulted in the up-regulation of X-linked genes. In conclusion, our findings describe a novel function for *C. elegans* condensin I/I^DC^ in organizing holocentric interphase chromosomes, which substitutes for the role played by cohesin in mammals.

## Introduction

The organization of eukaryotic genomes within the interphase nucleus is a complex process that involves extensive folding while maintaining essential functions such as replication, transcription, and repair. Chromatin conformation capture technologies and high-resolution imaging have been used to monitor genome conformation of interphase chromosomes^1–4^. In multicellular eukaryotes, the genome is organized across different scales. Long-range contacts between large megabase domains of two different types, A and B, create two compartments that can be identified as eu- and heterochromatin, respectively^2^. Over shorter distances, chromosomes are folded into loops and insulated 3D domains called topologically associated domains (TADs). TADs serve both to restrict the action of enhancers to specific promoters found within the same domain and to increase enhancer-promoter contact probability, resulting in co-regulation of genes within one TAD^5–10^. While A/B compartments correlate with the epigenetic status of the underlying chromatin, the formation of TADs is a consequence of DNA loop extrusion activities of the structural maintenance of chromosomes (SMC) complexes^11–14^. *In vitro* and *in vivo* experiments have demonstrated that these large ring-like complexes can generate loops of DNA and/or chromatin^15–18^. During interphase in mammals, cohesin is the major SMC loop extruder that interacts with oriented sequence elements bound by transcription factors like CCCTC-Binding factor (CTCF) that define boundaries between TADs as well as specific loops within TADs^19–21^.

Condensin, an additional SMC complex, is present in many organisms, from yeast to mammals, and is essential for the compaction and mechanical rigidity of mitotic chromosomes^22^. In budding yeast, condensin plays a crucial role in general chromosome compaction^23,24^, while in fission yeast, it slightly expands chromosomes and drives long-range organization into domains^25–27^. In many species, including mammals and *Drosophila,* two variants of condensin are present, namely, condensin I and II. The presence of the condensin II SMC complex correlates with 3D chromosome organization. Species without condensin II tend to have clustered centromeres and/or heterochromatin, while species with condensin II display chromosomal territories and poor centromeric clustering^28^. The effect of condensin II on chromosome structure results from lengthwise chromosome compaction during mitosis, which is retained in subsequent interphases due to the slow dynamics of long polymers such as chromosomes. In mammals, condensin complexes are either excluded from the nucleus (condensin I) or fail to bind chromatin outside of mitosis (condensin II)^29,30^. Notably, in *C. elegans,* at least one condensin complex is nuclear and active during interphase, an exception to the general rule of lack of condensin activity inside the nucleus during interphase.

Like most metazoans, worms express their homologs of cohesin, condensin I and II in somatic cells^31,32^. Two variants of the canonical cohesin complex are present, which can be distinguished by their kleisin subunits, COH-1 and COH-2/SCC-1^33^ (Fig. 1a). Analysis of RNAi phenotypes and localization of the two kleisins suggest a mitotic function for cohesin^SCC-1^ and non-mitotic functions for cohesin^COH-1^ ^32^. This early study demonstrated a differential localization of the two kleisins: SCC-1 is exclusively expressed in mitotic cells, while COH-1 is present in most nuclei^32^. In the absence of cohesin, many cells undergo catastrophic mitosis during early embryonic development^32,34,35^. Additionally, worms lacking cohesin subunits or the cohesin “loader” SCC−2/−4 (PQN-85/MAU-2) show defects in rDNA compaction in embryos^36^. However, the non-mitotic functions of both cohesins during interphase remain unclear, as little is known about cohesin localization on chromatin, and few non-mitotic phenotypes have been described to date^32^.

**Figure 1.**
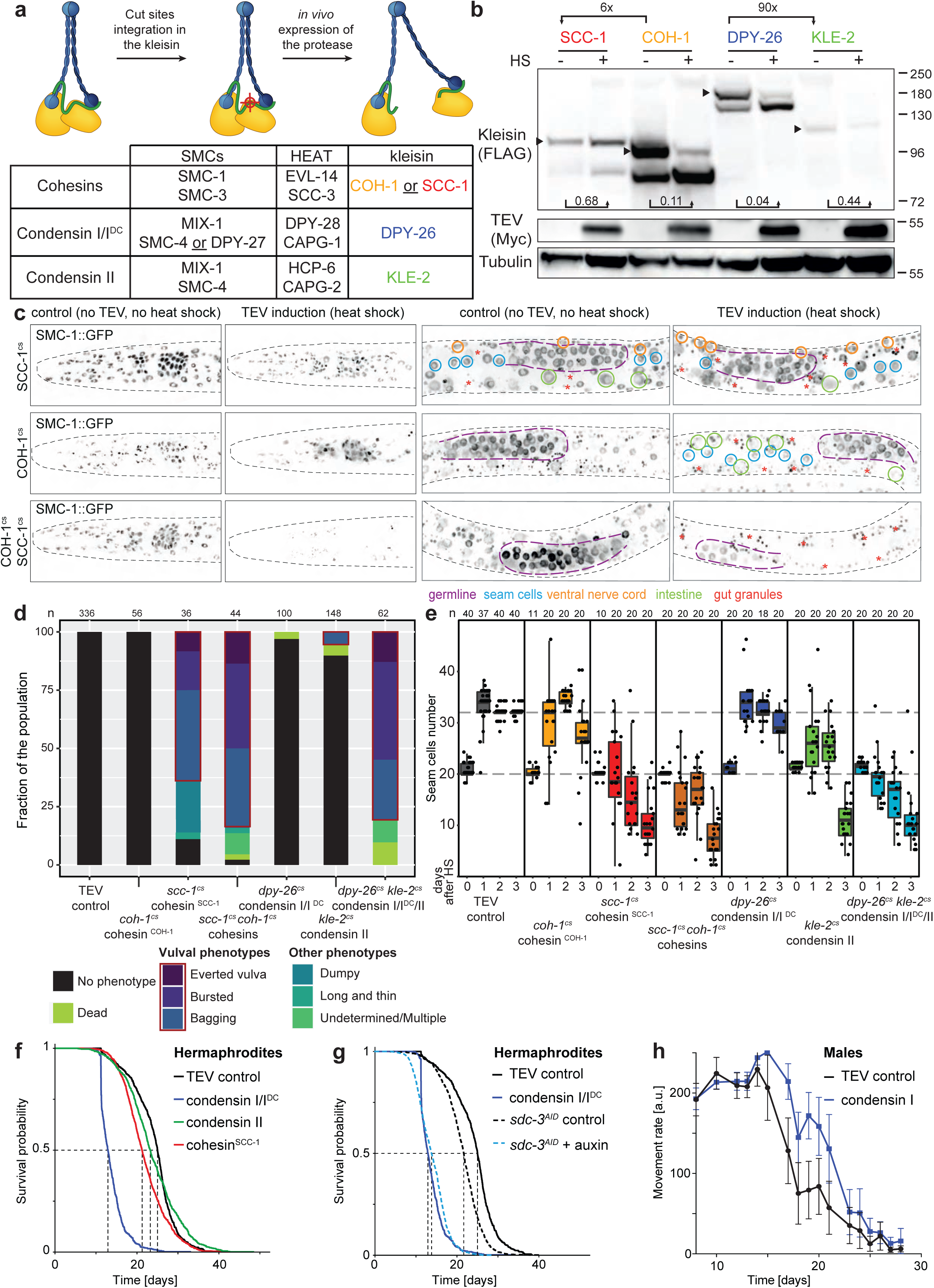
Loss of cohesin^SCC-1^ and condensin II causes disruption in post-embryonic vulval development and seam cell progression. **a.** Experimental approach for the functional analysis of SMC complexes *in vivo*. Adapted from ref. 33. SMC complex protein composition for condensin and cohesin is described in the table, with a color code for kleisin which is maintained throughout all figures. **b.** Comparison of individual kleisin expression levels and cleavage efficiency upon TEV protease expression. Quantification from at least 2 experiments (Fig. S1a). 3xFLAG-tagged kleisin expression was normalized to tubulin. **c.** Fluorescence signal in third larval stage animals expressing SMC-1::GFP, in the head section (left) or the middle section of the animals (right). The lighter hole in each nucleus corresponds to the nucleolus. For each section, the left panel corresponds to control animals (no heat shock, therefore no TEV protease expression), while the right panel shows animals in which TEV cleavage has been induced by heat-shock. Each line of images shows animals with TEV cleavage sites in SCC-1, COH-1 or both (top to bottom). For the middle section, cell lineage is indicated by circles and autofluorescence of gut granules by stars. **d.** Phenotypic outcome 72 hours post-induction of individual kleisin cleavage for the indicated SMC complexes (condensin I/I^DC^, condensin II, cohesin^SCC-1^ and cohesin^COH-1^) and multiple SMC complexes (cohesin^SCC-1^ ^and^ ^COH-1^ and condensin I/I^DC^/II). TEV control corresponds to a strain in which only the protease is expressed without any target cut site insertion. Color code below. **e.** Seam cells count per animal scored each day after kleisin cleavage induction of indicated SMC complexes. Color code as in A. **f.** Lifespan analysis of hermaphrodite animals after cleavage of the different indicated SMC complexes. Animals were transferred on day 1 of adulthood to plates with FUdR and scored automatically in the lifespan machine^108^. **g.** Lifespan analysis of hermaphrodite animals upon auxin-mediated degradation of the condensin I^DC^ loader SDC-3, comparing animals without auxin (no degradation) and with auxin (SDC-3 degradation). TEV control and condensin I/I^DC^ cleavage as in d. for the sake of comparison. **h.** Healthspan analysis of male animals in liquid medium upon condensin I cleavage using a wMicroTrackerONE device (male animals naturally crawl off Petri dishes, which makes lifespan analysis difficult on plates). Error bars from three biological replicates.

Nematode condensin I and II are essential for chromosome condensation during mitosis^31,37,38^. Deletions or RNAi knock-downs of many subunits of both condensins result in mitotic catastrophe in early embryos and nuclear defects in post-embryonically lineages^39–41^. A third condensin complex, condensin I^DC^, is also present and localizes specifically to the X chromosome in hermaphrodite animals during interphase. In this complex, the κ-SMC subunit SMC-4 of condensin I is replaced by its DPY-27 variant. The Sex Determination and Dosage Compensation complex (SDC) and a sequence motif called MEX enriched on the X chromosome specifically target condensin I^DC^ to the X chromosome^42^. Binding of condensin I^DC^ in hermaphrodite animals leads to down-regulation by half of X-linked gene transcription, compensating for the chromosomal imbalance between X and autosomes. Structurally, loading of condensin I^DC^ leads to the formation of TADs and loops that are unique to the X chromosome, indicating that the condensin I^DC^ variant is a *bona fide* loop extrusion factor^43–45^.

Here, we conducted a systematic investigation of the role of the four different SMC complexes in holocentric chromosome structure during somatic interphase by inducing cleavage of these complexes in larval stage worms, 90% post-mitotic. We performed phenotypic analysis, chromatin conformation capture, and RNA-seq to study the effects of SMC cleavage. Our results reveal that in contrast to mammals, cohesins play only a minor role in large-scale genome folding, while condensin I/I^DC^ is the primary loop extruder in nematodes. Cleavage of condensin I/I^DC^ reinforces the epigenetic-driven compartmentation on all chromosomes and results in the loss of TADs on the X chromosomes, similar to cohesin loss-of-function in mammals. Surprisingly, cleavage of condensin I/I^DC^ also leads to the appearance of an X-specific compartment enriched for the SDC complex. At the transcriptional level, we observe that cleavage of each individual SMC complex leads to moderate changes in gene expression on autosomes. However, condensin I/I^DC^ cleavage results in a major upregulation of X-linked genes and reduced animal lifespan due to its function in dosage compensation. In summary, our study highlights an unexpected function of condensin I isoforms in interphase genome folding in holocentric nematodes, with a major role in compartmentation regulation and dosage compensation on the X chromosome, while cohesins play only a minor role.

## Results

To interrogate the functions of individual SMC complexes, we constructed strains in which kleisin subunits are modified using genome editing to include three cleavage sites for the Tobacco Etch Virus (TEV) protease preceded by a FLAG tag (Fig. 1a)^46^. We created cleavable COH-1 and SCC-1 kleisins for the two somatic cohesins, and cleavable DPY-26 and KLE-2 kleisins for condensin I/I^DC^ and condensin II, respectively. Under non-heat-shock conditions, animals tolerated TEV cut sites in kleisins without exhibiting any apparent phenotypes. As a single FLAG tag was insufficient for kleisin detection by western blot, we added two additional tags to enable cleavage detection in otherwise identical strains. All kleisins were detected in third larval stage animals lysates using these triple FLAG strains (Fig. 1b) with COH-1, SCC-1 and DPY-26 being readily identifiable while KLE-2 signal was faint, likely reflecting its lower expression at this stage or expression in a subset of cells. For cohesin kleisins, COH-1 was 6 times more abundant than SCC-1, suggesting that most cohesin complexes are cohesin^COH-1^. This difference most likely reflects the expression of SCC-1 being limited to dividing cells^32^. Condensin I/I^DC^ kleisin DPY-26 was 90 times more abundant than condensin II kleisin KLE-2, suggesting that condensin I/I^DC^ is vastly predominant compared to condensin II (Fig. S1a). In the absence of induction and despite no detectable TEV protease, a small proportion of COH-1 and DPY-26 were cleaved in control conditions. This had no discernible effect on the animals as no phenotypes were identified in both the single or triple FLAG tag strains in the absence of TEV induction. Such phenotypes would have been expected, especially for DPY-26, due to its involvement in dosage compensation, whose absence leads to the appearance of short and fat (dumpy) animals. Upon TEV protease expression, COH-1 and DPY-26 were nearly completely cleaved (11% and 4% of uncut band remaining, respectively), while SCC-1 and KLE-2 full-length levels dropped by 32% or 50%, respectively (Fig. 1b, S1a). This ineffective cleavage of SCC-1 and KLE-2 might be due to a high turnover of these proteins and their complexes due to their mitotic functions. In particular, SCC-1 is cleaved by separase during the transition from metaphase to anaphase^22^, while degradation of *Drosophila* condensin II subunits CAP-H2 is necessary for full chromosome decompaction during interphase^47,48^. Throughout all experiments described below, we employed strains with a single FLAG tag.

Previous studies have shown that the two cohesin isoforms with COH-1 or SCC-1 are found to be expressed differentially^32^: COH-1 is expressed in most cells, SCC-1 is only present in mitotically active cells. To evaluate the cell-type specificity and effect of kleisin cleavage on both cohesin complexes, we analyzed the distribution and expression levels of SMC-1, one of the two SMC proteins present in both cohesin complexes, using an endogenously GFP-tagged SMC-1 protein^49^. Our findings show that SMC-1 was ubiquitously expressed at high levels in somatic cells, indicating that cohesins are present in most, if not all, somatic cells (Fig. 1c, controls). Additionally, SMC-1 was also present in the germline, where it forms a complex with three alternative kleisins COH-3, COH-4, and REC-8, which are exclusively expressed in this tissue (Fig. 1c, purple-circled region).

In our control experiments (no TEV induction), SMC-1::GFP fluorescence remained comparable in the presence of the cleavage sites in SCC-1, COH-1, or both kleisins. When SCC-1 was cleaved, we observed a minor change in overall fluorescence in somatic cells. However, when both COH-1 and SCC-1 were cleaved simultaneously, SMC-1 fluorescence disappeared completely in the soma, while germline nuclei retained their fluorescence. This finding indicates that simultaneous cleavage of COH-1 and SCC-1 not only opens the cohesin rings but also leads to the full degradation of the SMC-1 subunit. Moreover, when COH-1 alone was cleaved, many somatic nuclei lost fluorescence, consistent with COH-1 broader distribution^32^. The remaining fluorescent nuclei were either large, located among intestinal autofluorescence or aligned on the top of the animal’s bodies, indicating that they belonged to the intestinal or seam lineages, two tissues that are mitotically active in mostly post-mitotic animals (Fig. 1c, green and blue circled nuclei, respectively). Similar results were obtained when looking at the head of the animals where most nuclei retained fluorescence after SCC-1 cleavage, and all nuclei lost fluorescence when both SCC-1 and COH-1 were cleaved (Fig. 1c). When only COH-1 is cleaved, very few nuclei retain fluorescence, and while individual nuclei are hard to identify in the head, it is likely that these correspond to the handful of head neurons that are born post-embryonically. Overall, our experiments demonstrate that simultaneous cleavage of COH-1 and SCC-1 kleisins leads to the complete degradation of the SMC-1 subunit of the cohesin complexes in somatic cells. Furthermore, COH-1 is the most abundant cohesin kleisin, and its cleavage results in the complete disappearance of SMC-1 from most cells, except mitotically active ones.

### Mitotic functions of cohesins and condensins

To investigate the mitotic functions of SMC complexes, we induced kleisin cleavage during the first larval stage and assessed the resulting phenotypic consequences in subsequent development (Fig. 1d). At the time of induction, ninety percent of cells were post-mitotic, while the remaining cells were restricted to the hypodermal/seam lineages, the somatic gonad, and ventral hypodermal cells that give rise to the non-essential vulva, which allows egg-laying^50^. In wild-type worms, expression of TEV had no impact on development, while cleavage of cohesin^SCC-1^ resulted in developmental delays, morphological changes, and a range of phenotypes linked to a non-functional vulva in 64% of animals (Fig. 1d, S1c). Since the majority of cells were post-mitotic, this suggests that cohesin^SCC-1^ cleavage interferes with mitosis and/or terminal differentiation of the vulval precursor cells. In contrast, cleavage of cohesin^COH-1^ had no observable phenotype, and all animals eventually matured into phenotypically normal egg-laying adults. Simultaneous cleavage of both cohesin^SCC-1^ and cohesin^COH-1^ produced similar phenotypes as those observed for cohesin^SCC-1^, but with slightly higher penetrance, indicating minor mitotic roles for cohesin^COH-1^ and partial overlap of mitotic functions between the two cohesin variants.

In contrast to cohesin^SCC-1^ cleavage, individual cleavage of condensin I/I^DC^ or condensin II only slightly delayed late-stage development. Most worms (>90%) eventually matured into egg-laying adults without any other observable phenotype. However, simultaneous cleavage of both condensins resulted in the appearance of phenotypes associated with aberrant vulval development in a large majority of animals (Fig. 1d). Therefore, condensin I/I^DC^ and II can complement each other’s functions, but at least one type of condensin is necessary for faithful post-embryonic development.

To further investigate the roles of SMC complexes in mitosis, we examined seam cell divisions in L1 larvae following kleisin cleavage. These divisions can be monitored *in vivo* using specific fluorescent markers^51^. In this lineage, cell division is stereotypical and involves one asymmetric cell division per larval stage, resulting in one seam cell and one hypodermal cell that fuses with the rest of the hypoderm^50,52^. The number of seam cells remains constant after each mitosis, except for a symmetric division of a subset of the cells during the second larval stage, which increases the number from 20 to 32 cells per animal (Fig. 1e, TEV control animals).

Cleavage of cohesin^COH-1^ did not affect the number of seam cells, whereas cleavage of cohesin^SCC-1^ resulted in a decrease to less than 20 cells. Cleavage of both cohesin kleisins SCC-1 and COH-1 had an additive effect, causing a more severe decrease in seam cell numbers and misalignment of cells along the longitudinal axis of the body. The additive effect confirms that cohesin^SCC-1^ is the major mitotic cohesin in post-L1 larvae, and its loss alters both the integrity and orientation of the mitotic plane. Cleavage of condensin I/I^DC^ did not affect seam cell numbers, indicating that seam cell mitosis progressed faithfully in its absence. However, cleavage of condensin II resulted in a reduction in the number of seam cells to less than 20 cells within 48 hours, as well as misalignment of the mitotic axes (Fig. 1e). Cleavage of both condensins led to a more rapid decrease in seam cell number, reminiscent of the one observed upon cleavage of cohesin^SCC-1^ (Fig. 1e). Based on these findings, we conclude that, while quantitatively less abundant and with a lower TEV cleavage efficiency, condensin II and cohesin^SCC-1^ are necessary for post-embryonic cellular divisions, while condensin I/I^DC^ and cohesin^COH-1^ are dispensable.

Altogether, western blotting data, SMC-1 microscopy, and phenotypic analysis strongly suggest that even if the cleavage of some individual kleisin is not complete, the remaining levels of intact SMC complexes are not able to sustain their function. For both cohesins, one of the two main SMC subunits SMC-1 is degraded upon COH-1 and/or SCC-1 cleavage. For condensin I and II, complexes are not sufficient for faithful cell divisions. Although we cannot formally exclude that cleaved condensin kleisins might have dominant negative effects, the reviewmost likely explanation for the observations described above is that SMC complexes are no longer functional or degraded upon kleisin cleavage.

### Cleavage of condensin I/I^DC^, but not condensin II or cohesin^SCC-1^, drastically reduces lifespan

To assess the physiological effects of cleaving cohesin and condensin kleisins on the whole organism, we measured the lifespan of animals after cleavage. Cleaving condensin II or cohesin^SCC-1^ had little impact on lifespan, with animals living almost as long as TEV control animals (Fig. 1f). By contrast, cleaving condensin I/I^DC^ caused a drastic reduction in lifespan, with most animals dying by day 15 and only 3% still alive by day 20, compared to 71% of TEV control animals (Fig. 1f). As no mitotic phenotype was observed in animals after cleaving condensin I/I^DC^ alone, the reduction in lifespan is likely a consequence of the non-mitotic functions of this SMC complex.

### HiC analysis identifies genome organization at different scales

To better understand genome folding in post-mitotic nematodes, we conducted high-resolution chromosome conformation capture (HiC) on third larval stage animals, both in wild-type N2 and TEV control animals (Fig. 2ab, S2a, S3). As previously observed, the autosomes in control animals lacked the typical topologically associated domains (TAD) structure found in many metazoans and metaphytes, displaying instead three large contact domains (Fig. 2a): two multi-megabase domains located on the distal ends of the chromosomes (hereafter referred to as T domains, or telomere-proximal domains, corresponding to chromosomal arms) and a large domain located in the chromosome center (C domain). This organization correlates with the spatial segregation of repeat-rich heterochromatic T domains at the nuclear periphery and the transcriptionally active C domain located inside the nuclear lumen (for review, see ref. 53). Recently, this chromosome organization was described at the single-cell level in embryonic blastomeres using chromosome tracing techniques^54^. Perinuclear anchoring largely depends on H3K9 methylation marks enriched on chromosomal arms and the anchoring of these regions to the nuclear periphery via CEC-4, a chromodomain protein embedded in the nuclear envelope^55–57^. Genome-wide HiC matrices highlight this long-range organization within and between chromosomes (Fig. 3a-f, S2a). Additionally, a clear cross-hatched pattern is observed at a small scale (2-40 kb, Fig. 2a), both within and between chromosomes, indicating the presence of smaller compartments embedded into the large T and C domains and shared between chromosomes (Fig. 2a, S2a). Principal component analysis (PCA) on the genome-wide observed/expected contact matrix reflects this two-tiered organization (Fig. 2a bottom, S2a): the first eigenvector delineates large scale T/C domains, while the second captures the strong intra-chromosomal and weaker inter-chromosomal small-scale cross-hatch pattern (Fig. 2a, S2a).

**Figure 2.**
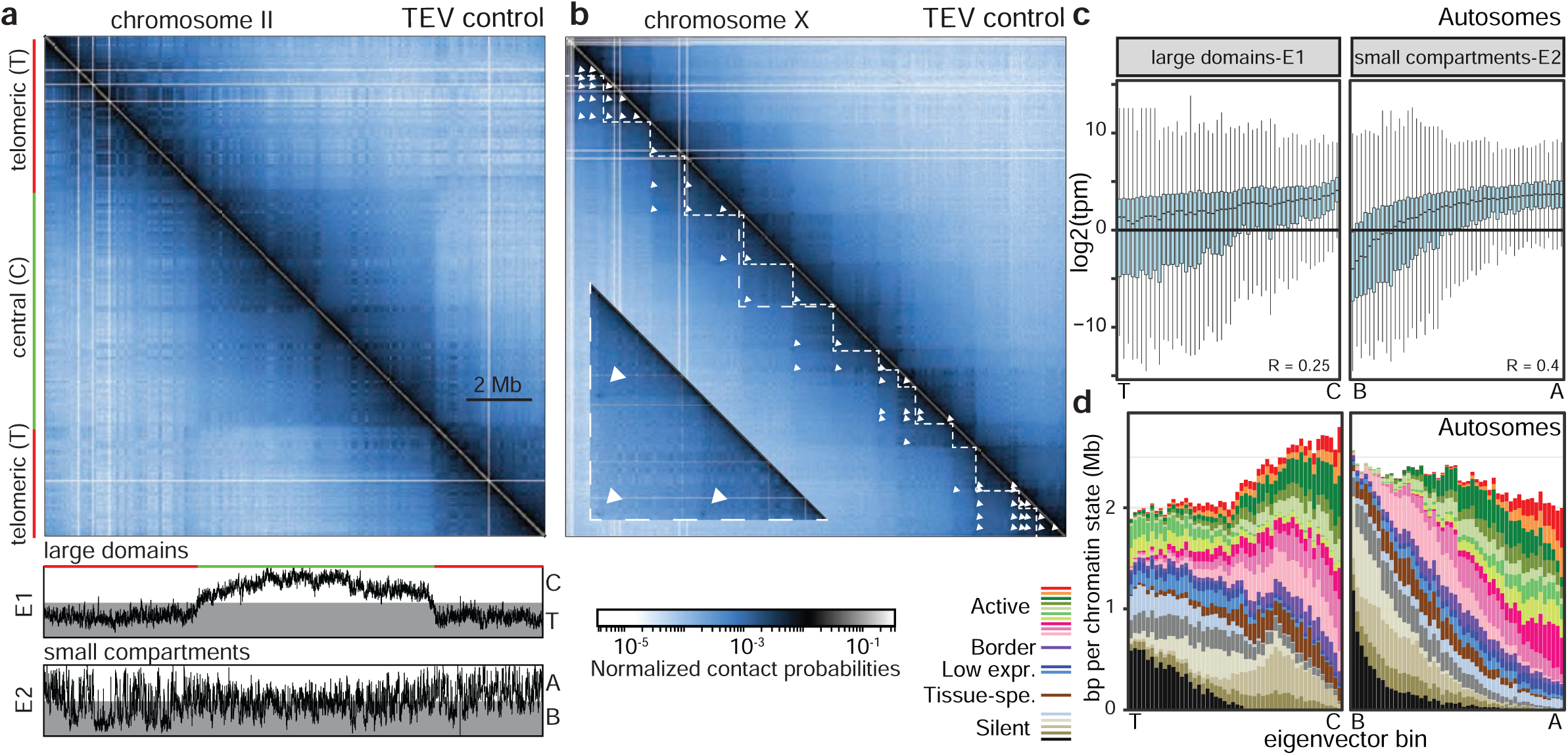
*C.elegans* interphase genome is organized at two different scales. **a.** HiC contact frequency map for chromosome II in TEV control animals. The compartment tracks at the bottom show the first and second inter-chromosomal eigenvectors of the PCA of the genome-wide matrix, with large, megabases size telomeric and central (T/C) domains identified in the first one, and small kb-size A/B compartments identified on the second one. T/C domains are delineated on the left side of the matrix. **b.** HiC contact map of chromosome X in TEV control animals. TADs and loops are highlighted in the lower part of the matrix (dashed lines and white arrowheads, respectively). A blow-up of the central part of the chromosome is shown superimposed on the lower part of the matrix (long dashed lines), highlighting the loops (white arrowheads). For both HiC maps, the thin white lines are unmappable genomic regions, for which no HiC contacts can be determined using short-read sequencing. **c.** Correlation of first (E1, large domains) and second (E2, smaller compartments) eigenvector values with transcription in TEV control animals. Eigenvector values for 2 kb genomic regions were binned into 50 equally spaced intervals. Genomic intervals in bin 1 have, therefore, the lowest 2% eigenvector values, while genomic intervals in bin 50 have the highest eigenvector values. For each 2 kb genomic region, the average expression values of the corresponding genomic intervals were determined and grouped into the 50 eigenvector value bins as a boxplot. Spearman correlation of log2 RNA-seq tpm *vs* eigenvalues for individual 2 kb genomic regions, is shown at the bottom right. **d.** Autosomal chromatin state composition of first and second eigenvector value bins. The chromatin state composition of the genomic intervals of each eigenvector value bin was determined using data from ref. 58. The correlation for autosomes is shown between chromatin states composition and eigenvector value for large T/C domains (first eigenvector) and small A/B compartments (second eigenvector). Similar results for the X chromosome, as well as the legend for chromatin states, are described in Fig. S2.

**Figure 3.**
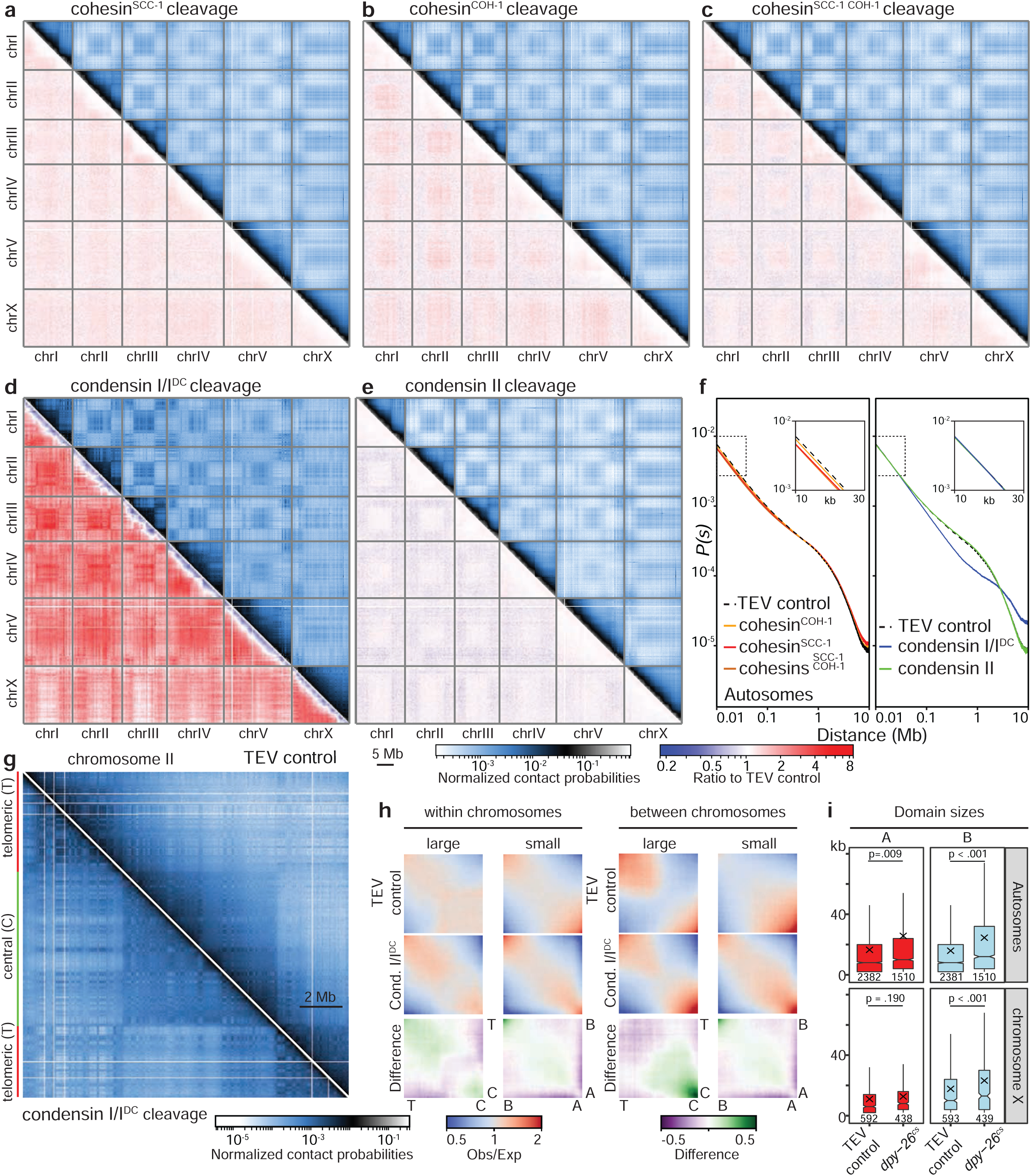
Condensin I folds the nematode genome. **a-e.** Genome-wide HiC contact maps in L3 animals upon SMC cleavage (upper right) and ratio to TEV control (lower left). The cleaved SMC complexes are indicated on top of the figure. Pooled data from two biological replicates per condition. **f.** Contact probability decay plots on autosomes for the different cohesin and condensin cleavages. Inset: contact probability decay at short distances (10-30 kb). **g.** HiC contact map of chromosome II in TEV control animals (upper right) and upon condensin I/I^DC^ cleavage (lower left), highlighting the increased A/B compartmentation. **h.** Saddle contact probability plots for large T/C domains and small A/B compartments within chromosomes (left) and between chromosomes (right) in TEV control (upper row), and in animals after condensin I/I^DC^ cleavage (middle row). Difference between the two conditions is shown in the bottom row. **i.** A and B compartment sizes on autosomes and chromosome X in TEV control and upon condensin I/I^DC^ cleavage (*dpy-26^cs^)*. The cross and the horizontal midline of the box denote the mean and median domain sizes, respectively, the box outlines values between the first and third quartiles.

While both the large domains and small compartments correlate with transcription, this correlation is far stronger for the small compartments (Fig. 2c; see methods for binning approach). A comparison of eigenvector values with previously characterized chromatin states, determined using chromHMM, confirmed this correlation with transcriptional activity (Fig. 2d, S2c, ref. 58). T domains are enriched for H3K9 methylated chromatin, but a significant portion also bears active histone marks, as transcriptionally active genes within T domains are interspersed among silent repeats. Conversely, C domains are enriched for active states, yet also harbor many Polycomb-marked regions (Fig. 2d, S2b). In contrast, small compartments faithfully separate active and silent chromatin: small compartments with high second eigenvector values are highly enriched for active chromatin states, while those with low values harbor inactive states, including both H3K9 and H3K27 methylation. These small compartments are, therefore, similar to A and B-type chromatin compartments previously described in mammalian systems, although a hundred times smaller in size ^2^.

Similar small A and B-type compartments have been described in nematodes using DNaseI HiC (ARC-C) and have been suggested to represent a TAD-like structure^59^. Indeed, in our HiC experiments, average contact maps on large A/B compartments (>10 kb) highlighted insulated domains (Fig. S2c), and the aggregated insulation score between A and B domains showed a minimum at the boundary (Fig. S2d). Our analysis confirms that these small compartments correlate with epigenomic states, as previously described in mammalian cells and *Drosophila*^2,60^. Interestingly, our findings reveal that the regions labeled as “mixed, tissue-specific” are distributed across all A/B small compartments eigenvalues (Fig. 2d). This is in contrast to the active and silent chromatin states, which are typically associated with specific compartments. Since HiC experiments are conducted on whole animals that contain a diversity of tissue-specific genes, one would expect tissue-specific genes to join the A or B compartments depending on their cell type-specific chromatin state^61^, while the global genome folding into T and C domains is conserved across all cell types.

In hermaphrodite worms, the X chromosomes fold into TADs, setting them apart from autosomes. TADs are formed by the X-specific activity of condensin I^DC^. This phenomenon has been extensively studied and documented^43,45,62,63^. While most of the X chromosome lies within a large C domain (data not shown), small compartments within the TADs form a cross-hatched pattern that, similar to autosomes, correlates with chromatin states (see Fig. S2b). Moreover, the presence of high contact probability signals at the corners of the TADs suggests the existence of more stable loops between TAD boundaries, which have been observed in HiChIP experiments with the condensin I^DC^ subunit DPY-27^44^. These loops can span over two or more of the TAD boundary elements that are defined at the sequence level by the strong Motif Enriched on X (MEX) motifs^64^. The white arrows in Fig. 2b indicate the location of these stable loops. The unique folding and looping pattern of the X chromosome TADs are key features that distinguish them from autosomes.

### Condensin I/I^DC^ is the major SMC complex for long-range interphase genome folding

In mammalian cells, cohesin is involved in interphase chromosome folding besides its role in mitosis^11–13^. Cohesin extrudes DNA loops, and this activity is blocked at oriented boundary sequence elements recognized by the Zn-finger DNA binding factor CTCF^16,20,21^. While the role of cohesin in mammalian interphase chromosome folding has been extensively studied, the identity of the SMC complex that drives chromatin looping in interphase autosomes in nematodes remains unknown. To address this question, we performed Hi-C experiments following the cleavage of individual cohesin or condensin kleisins, as well as the simultaneous cleavage of both cohesin kleisins. In contrast to mammalian cells, in which cohesin depletion or ablation leads to a sharp decrease in contact probabilities reflecting the loss of long-range looping, we observed that cleavage of either of the two worm cohesins individually or both cohesins together only decreased contact probabilities to a small extent at short distances (10-100 kb, Fig. 3abc). We observed a slightly larger decrease in contact probabilities upon cleavage of cohesin^SCC-1^ compared to cohesin^COH-1^ (Fig. 3f). Simultaneous cleavage of both cohesins led to an additive, yet a very minor decrease of the short-range contact probabilities (Fig. 3f). It is possible that the small decrease in contact probabilities observed for cohesin^SCC-1^ arises from the small fraction of mitotic cells where it is expressed since Hi-C experiments were performed on whole animals.

Taken together, our findings suggest that nematode cohesins, unlike their mammalian counterparts, have no significant impact on large-scale interphase chromosome folding. Similar to cohesins, condensin II kleisin cleavage did not lead to significant changes in HiC maps, indicating that its primary function is to ensure proper mitotic chromosome compaction (Fig. 3ef). In stark contrast, cleavage of condensin I/I^DC^ led to significant changes in contact frequency maps. Specifically, we observed a decrease in contact probabilities as compared to control experiments at distances larger than 100 kb, suggesting that condensin I/I^DC^ is involved in promoting these long-range contacts. Interestingly, for distances over 3 Mb, the contact probability was higher after condensin I/I^DC^ cleavage than in control experiments or cleavage of other SMC complexes (Fig. 3df), indicating that condensin I/I^DC^ actively represses the formation of these very long-range contacts. Taken together, our results support the hypothesis that condensin I/I^DC^ is the loop extrusion factor responsible for creating long-range interphasic chromatin loops in nematodes.

### Condensin I cleavage increases epigenomic compartmentation

By analogy with observations made in mammalian cells upon depletion of cohesin, one would predict that a consequence of condensin I/I^DC^ cleavage would be the reinforcement of A and B compartments^11–13^. Indeed, condensin I/I^DC^ cleavage, but no other SMC cleavage led to an increase of contact frequency in the checkerboard and cross-hatch patterns (Fig. 3g, compare TEV control to condensin I/I^DC^ cleavage). This resulted in the reinforcement of T and C domains within chromosomes, as well as B compartments (quantified in Fig. 3h). Moreover, the contact probability between autosomal central C domains and between B compartments increased significantly between chromosomes (Fig. 3h). A and B compartments also saw an average size increase of 62% and 54%, respectively (Fig. 3i). Notably, chromosome X showed an increase in inter-chromosomal contact probability with the T domains of each autosome (Fig. 3d), suggesting a relocation of the X chromosome towards the nuclear periphery upon condensin I/IDC cleavage, as previously observed upon loss of dosage compensation^65^.

To determine whether genomic regions that switch compartments have specific genomic or epigenomic features, we divided the T/C and A/B compartment tracks (E1 and E2 eigenvectors, Fig. 2a) into equally-sized groups or terciles (high (H), middle (M), and low (L) eigenvector values). We then monitored 10 kb bins of the genome that switched from one tercile to another upon kleisin cleavage (Fig. S4). Very few regions shifted more than one tercile upon cleavage for any kleisins. However, a significantly higher number of changes were observed upon condensin I/I^DC^ cleavage, with 21% of genomic bins switching terciles (compared to 8-12% for other kleisins). Moreover, we found that condensin I/I^DC^ E2 bins switching down from H to M were depleted in annotated transposable elements but enriched in non-coding RNAs and tRNAs, active histone marks, and harbored genes with an increase in expression upon cleavage. In contrast, bins switching up from M to H or L to M did not show such trends (Fig. S4).

Our analysis leads us to conclude that the primary SMC complex responsible for chromosome folding in *C. elegans* is condensin I/I^DC^, whereas cohesins are mainly involved in short-range looping. Importantly, cleavage of condensin I/I^DC^ led to genome decompaction and an increase in inter-chromosomal contacts. Additionally, we observed a significant enhancement of T/C and A/B compartmentation, which provides further insight into the role of this complex in the three-dimensional organization of the genome.

### Condensin I/I^DC^ cleavage leads to X chromosome TAD disappearance and reinforcement of an X-specific loop compartment

In mammalian cells, loss of cohesin leads to the disappearance of TADs and loops^11–13^. Since these structures are only present on the X chromosome in *C. elegans*, we investigated how cleavage of the different SMC complexes affects them. Cleavage of either cohesin kleisins resulted in a slight loss of contact probability at short distances (10-100 kb), while condensin II kleisin cleavage had no effect (Fig. S2h). In contrast, condensin I/I^DC^ cleavage caused major changes on a large scale (>100 kb, Fig. 4a, compare TEV control and condensin I/I^DC^), with X-specific TADs disappearing and insulation between TADs being lost or greatly weakened (Fig. 4abd). However, cleavage of the putative loop extruder condensin I/I^DC^ did not lead to the loss of the high contact frequency spots located at the TAD corners. Instead, it resulted in the formation of reciprocal loops between thirty-five small regions, creating dotted lines on the Hi-C contact probability map (Fig. 4a). Manual mapping and automatic detection^66^ of the loops or their anchors showed a high overlap between loop anchors and TAD boundaries present in wild-type and TEV control animals(Fig. 4ab).

**Figure 4.**
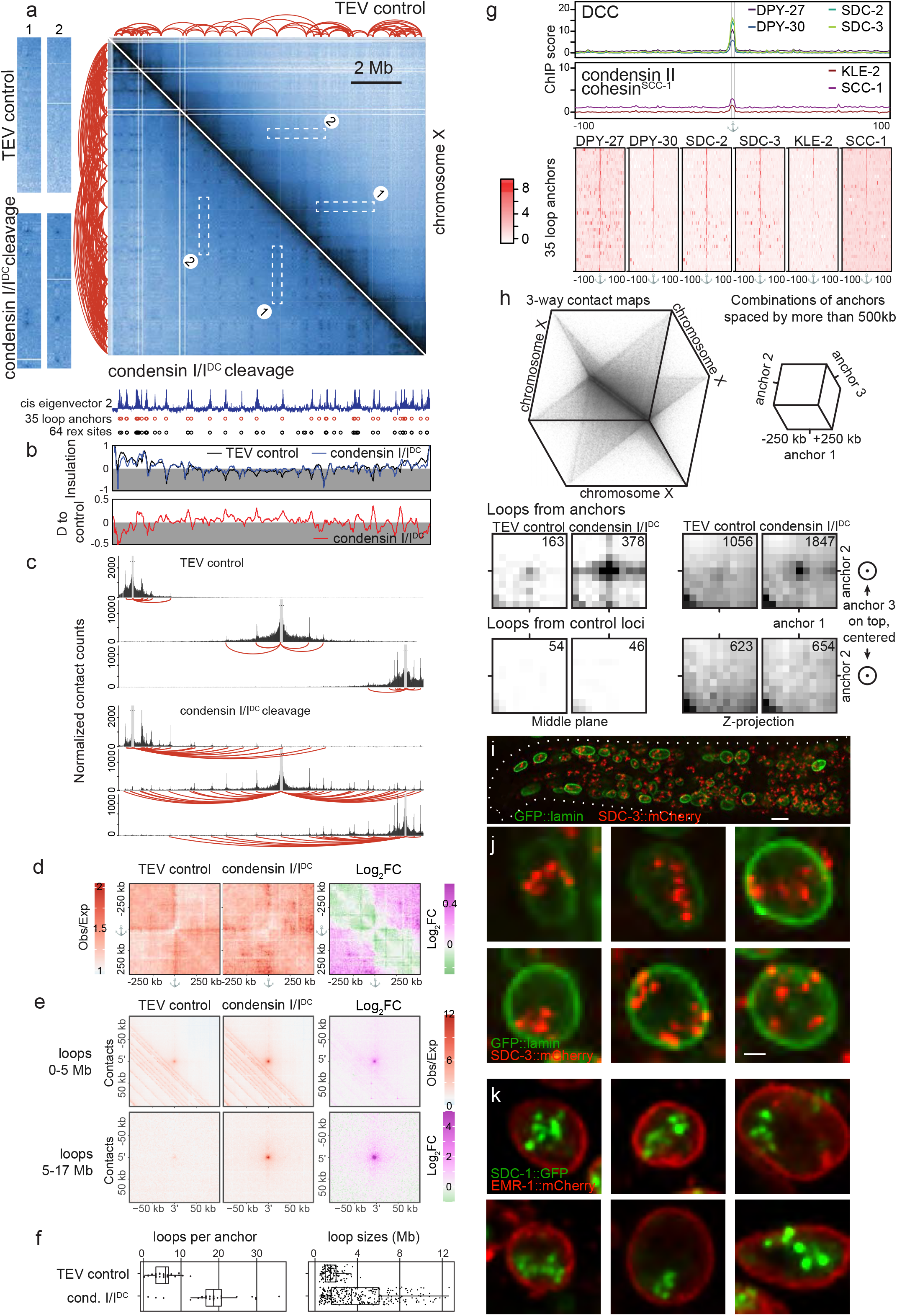
The hermaphrodite X chromosomes form a rosette-like structure. **a.** HiC contact frequency matrices in control (upper right) and upon cleavage of condensin I/I^DC^ (lower left). Left side: comparison of regions depicted in the map on the left highlighting the formation of dotted lines of high contact probabilities upon condensin I/I^DC^ cleavage. Chromatin loops are depicted as arches on the top (TEV control) and the left side (condensin I/I^DC^ cleavage). Bottom: second *cis* eigenvector for condensin I/I^DC^, used for loop anchor detection (local maxima), corresponding loop anchors and *rex* sites^64^. **b.** Insulation scores for the two matrices in 4A and the difference between them. **c.** Virtual 4C analysis on 3 representative loop anchors in the TEV control and upon condensin I/I^DC^ cleavage. Loops are depicted as arches. **d.** Average observed-over-expected Hi-C contact maps centered on the anchors defined upon condensin I/I^DC^ cleavage, highlighting their correlation with TAD boundaries in control conditions and the loss of these boundaries upon condensin I/I^DC^ cleavage. **e.** Average observed-over-expected Hi-C contact maps centered on all possible loops between pairwise anchors combinations. Loops are segmented between short (<5Mb) and long-range (>5Mb) ones. Average HiC maps are shown in TEV control and upon condensin I/I^DC^ cleavage, as well as the log2 fold change (Log_2_FC) between the two conditions. **f.** Number of loops per anchor and loop sizes in TEV control and upon condensin I/I^DC^ cleavage. **g.** Average (top) and individual (bottom) ChIP-seq enrichment of the protein subunits of DCC (DPY-27 - L3 animals, DPY-30, SDC-2, SDC-3 - embryos), condensin II (KLE-2 - L3 animals) and cohesin (SCC-1 - L3 animals) for the 35 loop anchors detected upon condensin I/I^DC^ cleavage. Data from refs. ^67,75 64^ and ^59^. **h.** Detection of 3-way contacts between loop anchors identified in A. 3-way contact maps (top left) were built from HiC reads containing more than two fragments ligated together. Average 3D HiC maps between all combinations of three loop anchors spaced by at least 500 kb are shown at the bottom, either the middle plane (left) or the sum Z-projection (right), in the TEV control experiment and upon condensin I/I^DC^ cleavage. The number on the upper right corner is the number of identified 3-way contacts in the middle voxel/pixel. **i.** Nuclear pattern of SDC-3::mCherry (red) in animals in control conditions. GFP-lamin delineates the nuclear periphery. Bar: 5 μm. **j.** Detail of individual nuclei from the animal in I. Bar 1 μm. **k.** Nuclear pattern of SDC-1::GFP (green) in animals in control conditions. EMR-1::mCherry delineates the nuclear periphery (red).

While each anchor would create contacts with a limited number of neighboring boundaries in control situations (median 6; Fig. 4cf), cleavage of condensin I/I^DC^ led to an increase in the number of loops per anchor (median 19; Fig. 4cf). Additionally, the mean size of loops increased from 1.25 Mb in control animals to 3.7 Mb upon condensin I/I^DC^ cleavage, with many loops having sizes larger than 8 Mb, which represents half the entire X chromosome length (Fig. 4f). Average Hi-C contact frequency maps using all possible loops between pairwise anchors confirmed that contact frequencies between anchors increased both at small (<5 Mb; 30% increase relative to background) and large scale (5-17 Mb; 80% increase; Fig. 4e). MEME motif detection identified the MEX motif in 35 loop anchors, and 34 out of 35 co-localized with previously identified *rex* sites, a subset of MEX motifs with high enrichment for the X-specific condensin I^DC^ targeting complex (Fig. 4a, bottom, Supplementary Table 6)^64^.

Using a previous classification of rex sites as strong, intermediate, and weak^64^, we observed that 15 out of 17 (88%) strong *rex* sites overlap with loop anchors. Only half (8 out of 16) intermediate *rex* sites and 9 out of 31 (29%) weak *rex* sites overlapped with anchors, respectively. Further comparisons with previously published ChIP-seq data demonstrated strong enrichment at loop anchors for condensin I^DC^ and the X-specific targeting complex subunits SDC-2, SDC-3, and DPY-30 (Fig. 4g). Loop anchors were found to colocalize with a subset of the highest ChIP peaks for these proteins, but not all peaks, with an average enrichment between 44 and 157 times higher than the average enrichment in the surrounding 100 kb regions (Fig. 4g). Surprisingly, the other SMC complexes, cohesin^SCC-1^ and condensin II, showed a similar enrichment at loop anchors, suggesting that the anchors are boundary elements for all SMC complexes or loading sites for these as hinted earlier for condensin II (Fig. 4g)^67^.

We next investigated whether condensin I/I^DC^-independent loop anchors form pairs or clusters of more than two anchors. To do this, we analyzed HiC reads that had three or more loci ligated together (726’736 and 635’546 3-way contacts on the X chromosome for TEV control and condensin I/I^DC^ cleavage, representing 0.17% of all contacts for both libraries, respectively). Using these contacts, we constructed three-dimensional contact maps at 50kb resolution (Fig. 4h)^11,44,68^. Using those maps, we analyzed whether loop anchors form clusters, comparing all combinations of three anchors with more than 500 kb between them, as well as combinations between a control set of loci located 500 kb away from the anchors (Fig. 4h). In the TEV control libraries, we detected 3-way contacts between loop anchors, confirming that *rex* sites in loop anchors cluster together as previously reported^44^. However, upon condensin I/I^DC^ cleavage, the 3-way contact frequency between loop anchors increased more than two-fold, demonstrating that at least three loop anchors indeed come together. No significant 3-way contacts were observed for control loci in any condition. These findings suggest that condensin I/I^DC^-independent loop anchors tend to form clusters involving three or more loci, indicating a rosette-like structure of the X chromosome where anchors assemble in one or more hubs to which individual TADs are attached.

### The condensin I^DC^ loader complex protein SDC-3 and SDC-1 form nuclear bodies

We observed that loop anchors are highly enriched for the SDC complex, which is essential for the enrichment of condensin I^DC^ onto the X chromosomes^69^ (Fig. 4g). We, therefore, asked whether clustered loop anchors would be visible *in vivo* by adding fluorescent tags to the SDC-1 and SDC-3 proteins at their endogenous gene location. Consistent with previous studies^70–72^, the proteins are expressed exclusively in hermaphrodite animals, show diffuse localization during early embryonic development (less than 40 cells), and label one or two nuclear territories in animals after dosage compensation onset (data not shown and Fig. 4ij). Strikingly, SDC-1 and SDC-3 fluorescence was not homogenous inside the territory but clustered in individual spots that we named SDC nuclear bodies. We observed little to no background fluorescence outside of the clusters (Fig. 4ijk). Although linking ChIP, HiC, and microscopy data is challenging^73^, this strongly hints at the presence of loop anchor aggregates in the wild-type condition.

Interestingly, we found that SDC bodies fluorescence did not differ between the control and condensin I/I^DC^-cleaved conditions (Fig. S5), indicating that the maintenance of SDC bodies is independent of condensin I/I^DC^ activity and the presence of TADs. Assuming these spots represent clustered loop anchors, the absence of modification of the fluorescence pattern indicates that the rosette structure of the X chromosome is present in the control situation, yet not visible in HiC maps due to the high number of contacts present inside the TADs. Together, this suggests that the maintenance of SDC bodies is independent of condensin I/I^DC^ activity and the presence of TADs, while TADs themselves require condensin I/I^DC^ to persist.

### Cohesin^SCC-1^ and condensin II cleavage lead to correlated transcriptional changes

We next examined the effects of SMC cleavage on gene expression by comparing animals after cleavage to TEV control animals. Our analysis showed that with the exception of X-linked genes after cleavage of condensin I/I^DC^, SMC cleavage had modest effects on gene expression (log2 fold change (LFC) between 0.15 and 0.5, Fig. S6a, Supplementary File 1). Exon-Intron split analysis^74^ confirmed that most changes occurred at the transcriptional level (Fig. S8). Considering genes as significantly differentially expressed when |LFC|>0.5, we found that only 3-12% of genes show different expression levels (154 to 1509 genes, Fig. 5a). Upon cleavage of condensin II, cohesin^COH-1^, cohesin^SCC-1^ or cohesin^COH-1/SCC-1^, these genes were distributed evenly across all chromosomes, with slightly more genes upregulated than downregulated. This suggests a repressive function for SMC complexes in gene regulation (Fig. 5a, S6c). Cleavage of both cohesin^SCC-1^ and cohesin^COH-1^ kleisins led to a synergistic effect, with the highest number of misregulated genes (Fig. 5ab, S6c).

**Figure 5.**
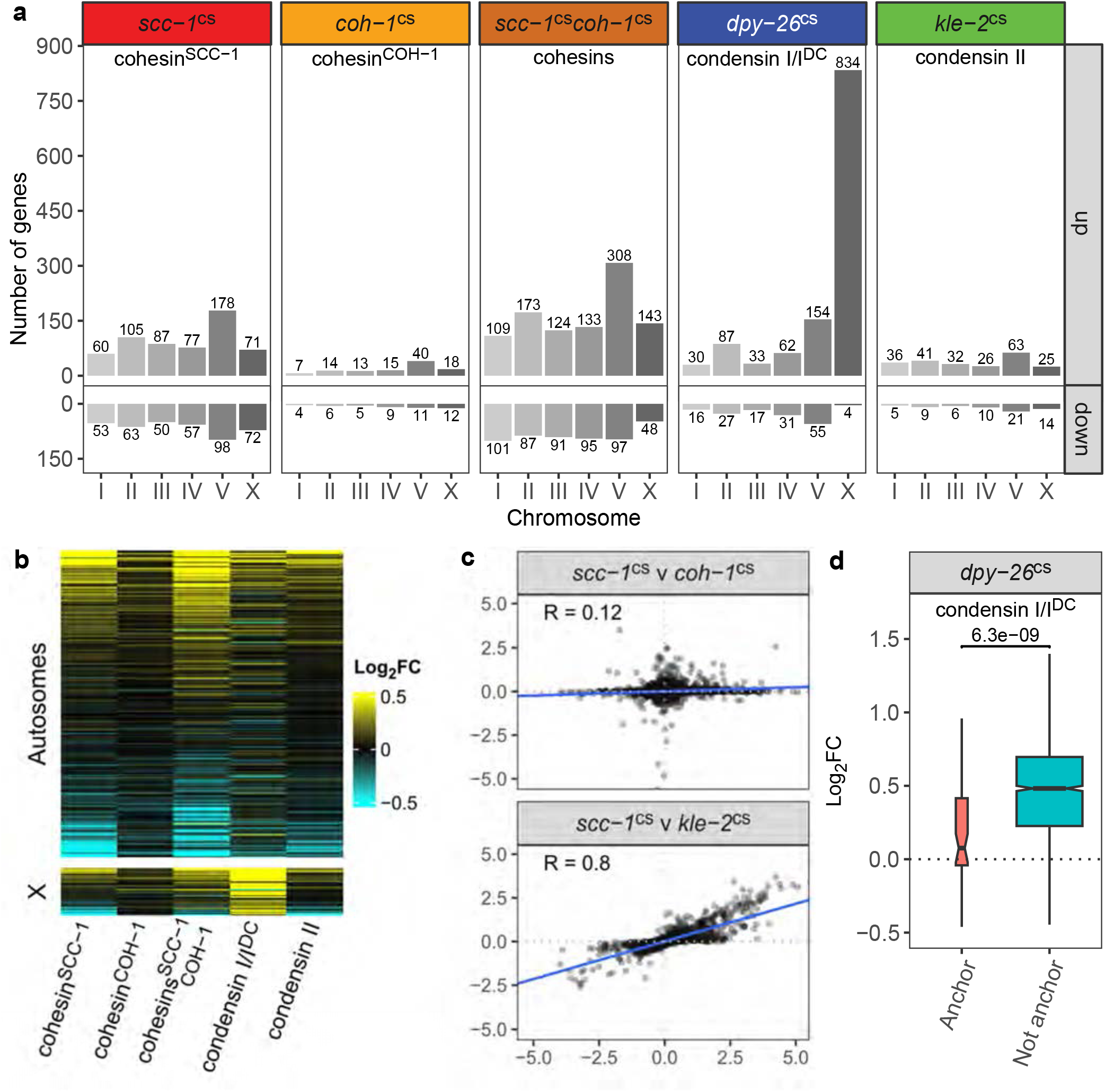
Differential gene expression after cleavage of kleisin subunits of different SMC complexes. **a.** Number of significantly up and down-regulated genes (AdjP<0.05, |Log2 Fold Change (LFC)|>0.5) per chromosome upon cleavage of the alternative kleisin subunits of cohesin, *scc-1^cs^*and *coh-1^cs^* individually or together, or condensin I/I^DC^ (*dpy-26^cs^*) and condensin II (*kle-2^cs^*). The number of up/down regulated genes is shown next to each bar. **b.** Hierarchical clustering of the LFC of 12465 genes filtered by DESeq2 as expressed in at least one dataset. Top panel: 10792 autosomal genes. Bottom panel: 1673 X-linked genes. **c.** Correlation of the LFC of 13734 expressed genes (with at least 10 read counts in total) from the datasets of the two alternative cohesin kleisins (*scc-1*^cs^ and *coh-1*^cs^, top panel) and the datasets of condensin II and cohesin^SCC-1^ cleavage (bottom panel). The blue line indicates the best fit linear regression, and the Pearson correlation coefficient is shown. **d.** LFC after cleavage of condensin I/I^DC^ of 55 genes that lie within 10 kb of the 35 loop anchors defined in Fig. 4a (“Anchor”), compared with 1707 genes found on the X chromosome that do not overlap with loop anchors (“Not anchor”). The p-value from the Wilcoxon rank sum test comparing genes in anchor to non-anchor regions is shown in the panel.

We observed little overlap between differentially regulated genes upon individual cleavage of the two cohesin isoforms, and the correlation of their LFC was weak (Pearson’s r=0.12, Fig. 5c, top panel). This suggests that the two cohesin complexes have different functions or targets in gene regulation. In contrast, there was a clear correlation between differentially regulated genes upon cleavage of cohesin^SCC-1^ and condensin II (266 genes overlap, Pearson’s r=0.8, Fig. 5c, bottom panel, S7ab). Gene Ontology analysis revealed a significant proportion of the genes were linked to proteolysis or stress response, possibly in relation with the shared mitotic functions of cohesin^SCC-1^ and condensin II (Supplementary File 2). This stress response was not the result of the heat shock treatment used to activate the TEV protease, as an independent dataset of genes misregulated in *kle-2* null L3 animals^67^ similarly correlated well with our condensin II or cohesin^SCC-1^ cleavage datasets (Pearson correlation 0.22, 0.25, respectively). In summary, our results indicate that cleavage of either cohesins or condensin II have relatively minor transcriptional consequences, but a common set of genes appears to be regulated by cohesin^SCC-1^ and condensin II, likely due to their common mitotic function.

### Cleavage of condensin I/I^DC^ or knock-down of the X-targeting condensin I^DC^ complex mainly impacts genes on the X chromosome

The cleavage of condensin I/I^DC^ led to the most significant changes in gene expression on chromosome X. Half of the expressed X-linked genes showed significantly higher expression (834 out of 1604), while only 4 were downregulated (52% and 0.0025% of expressed X-linked genes, respectively, Fig. 5ab, S6bc). Therefore, the main effect of condensin I/I^DC^ cleavage on gene expression is the relief of repression of X-linked genes, due to the cleavage of condensin I^DC^ exclusively bound to the X chromosome^42^. A comparison with previous datasets showed a significant overlap between X-linked genes upregulated following condensin I/I^DC^ cleavage and those upregulated following condensin I^DC^ subunit *dpy-27* RNAi knock-down^75^ or mutation of the DCC-associated *dpy-21* H4K20 demethylase gene^62^ (25% and 46%, respectively, Fig. S10d). In contrast to mammalian cells upon cohesin depletion^12^, we found no evidence for pervasive upregulation of non-coding RNAs, nor for general misregulation of transposons or repetitive sequences for cohesin^SCC-1^, condensin II or condensin I/I^DC^ cleavage (data not shown). The same result was obtained when sequencing ribosomal-depleted total RNA instead of poly-adenylated transcripts.

Interestingly, X-linked gene upregulation in close vicinity to the loop anchors characterized above and bound by the SDC complex was significantly lower upon condensin I/I^DC^ cleavage (Fig. 5d, S7c). In non-anchor regions, 48% of genes (823 of 1707) were significantly upregulated, while in 10 kb regions around loop anchors, only 11% of genes (6 of 55) were upregulated. This finding indicates that gene regulation is largely maintained close to sites bound by the SDC subunits of the DCC but not inside the TADs delimited by these loop anchors.

### Lack of dosage compensation, not autosomal chromatin decompaction, leads to reduced lifespan

Our HiC and RNA-seq data reveal two outcomes of condensin I/I^DC^ cleavage: genome-wide loss of chromosome compaction on both autosomes and the X chromosome, and upregulation of X-linked genes primarily due to loss of dosage compensation. Either of these phenomena could cause the observed reduction in lifespan observed upon condensin I/I^DC^ cleavage (Fig. 1f). To distinguish between these two possibilities, we analyzed the lifespan of degron-tagged SDC-3 animals. SDC-3 is essential for condensin I^DC^ enrichment on the X chromosomes but not for condensin I loading onto autosomes^70^. Inducing SDC-3 degradation with auxin led to a decrease of the TAD structure of the X chromosomes, as well as upregulation of X-linked genes, albeit to a lesser degree than condensin I/I^DC^ cleavage (Fig. S9-10). When grown on media without auxin, animals with a degron tag on SDC-3 and expressing the auxin-binding receptor TIR1 had a lifespan similar to that of TEV control animals. The slightly longer lifespan of the latter was likely due to the heat-shock used to induce TEV expression, which was previously shown to increase lifespan^76^. However, lifespan analysis showed that upon SDC-3 degradation, most animals were dead by day 15, and only 4% of the animals remained alive by day 20, similar to condensin I/I^DC^ cleavage (Fig. 1g). Moreover, male animals, which do not require dosage compensation and do not express condensin I^DC^, showed a slight increase in lifespan upon condensin I cleavage (Fig. 1h). We conclude that the reduction in lifespan after condensin I/I^DC^ cleavage is not caused by genome-wide unfolding but by the X-specific disruption of chromosome structure and the consequent upregulation of X-linked genes.

## Discussion

In this study, we examined the function of SMC complexes cohesin^COH-1^, cohesin^SCC-1^, condensin I and condensin II in fully differentiated nematodes using an inducible cleavage system to open the SMC rings. Our findings indicate that cohesin^SCC-1^ is crucial for mitosis, consistent with its exclusive expression in mitotic cells ^32^. In contrast, cohesin^COH-1^ does not seem to play a role in mitosis, suggesting that the two cohesin variants have different, non-redundant functions. In contrast to RNAi studies against single subunits of condensin I/I^DC^ in embryos and in larval intestinal cells^31^, condensin I and II are individually dispensable for faithful cell division. This suggests that there may be different requirements for condensin function in different cell types and stages of development. The absence of phenotype of individual condensin cleavage in slowly dividing cells is reminiscent of the subtle changes observed in mammalian cells upon individual condensin subunit depletion^77,78^: in cultured cells, the depletion of condensin I or II subunits changes the mechanical rigidity of the mitotic chromatids and delays chromosome condensation during early prophase but does not impair mitotic progression. However, simultaneous cleavage of both condensin I and II in nematodes results in mitotic defects, which is consistent with what has been observed in mammalian cells.

Furthermore, we performed high-resolution chromosome conformation capture in wild-type control animals and revealed new insights into the folding of the nematode genome. Our findings suggest that two levels of organization direct genome folding. Firstly, consistent with previous studies, the outer thirds of the autosomal arms, which are rich in repeats and heterochromatin, form large telomeric (T) perinuclear domains^55,79–82^, while the central regions of autosomes, which are transcriptionally more active, are located in the nuclear lumen (large C domains). Secondly, we observed that small regions of the genome with similar transcriptional status and epigenomic marks preferentially interact with each other together, forming distinct small A and B compartments. These compartments have sizes ranging from 2 to 100 kilobases, which is 10 to 100 times smaller than the ones observed previously in mammalian and *Drosophila* genomes^2,83^.

In our study, we used individual SMC cleavage in combination with chromosome conformation capture to investigate the role of each SMC complex in chromosome folding in mostly post-mitotic animals. In mammalian cells, chromatin looping and megabase-scale TAD formation occur through the loop extrusion activity of cohesin coupled with boundary elements such as CTCF binding sites. Our findings demonstrate that in *C. elegans*, the main SMC complex involved in large-scale chromosome folding is condensin I/I^DC^ rather than cohesins, which have limited activities in short-range looping. Cleavage of condensin I/I^DC^ leads to both increased genome decompaction and inter-chromosomal contacts, as well as augmented T/C and A/B compartmentation. Similar long-range looping activity (>100 kb) of condensin has been described in budding and fission yeast^23,27^, and mutations in the condensin complex in these organisms also led to increased chromosomal mixing. Whether chromosomal mixing is a direct effect of condensin mutations remains unclear, as acute depletion of the SMC subunit Cut14 did not reproduce the mixing observed in the mutant^27,84^. We conclude that in nematodes, condensin I variants are responsible for both autosomes and chromosome X folding^43^.

Why would condensin I in worms take over the role of cohesin in mammals? In mammals, most cohesins are removed from chromosome arms right before anaphase, while centromeric complexes ensure sister chromatid cohesion^22^. However, the holocentric nature of the nematode chromosomes may make it difficult to differentiate between centromeric and non-centromeric cohesin populations, as worm centromeres are defined by the binding of transcription factors and active transcription^85^. Repurposing condensin I/I^DC^ as an alternative long-range loop extruder while duplicating cohesins with one variant - cohesin^SCC-1^ - exclusively expressed in dividing cells would functionally separate SMC complexes involved in sister chromatid cohesion from the ones involved in interphasic genome looping^32^.

Notably, nematodes lack a CTCF homolog and any well-characterized boundary element or factor that is involved in the formation of TAD boundaries in mammals or *Drosophila*. This suggests that genome folding in nematodes primarily relies on T/C domains and smaller epigenomic A/B compartments, rather than TADs^83,86^. The absence of TAD structure on autosomes implies that in nematodes, transcriptional regulation is mainly carried out by proximal enhancers that are located close to their target promoters, as the activity of enhancers would not be limited by TADs as in mammals^10^. In fact, all the enhancer sequences in a compendium that demonstrated transcriptional activation activity of minimal promoter reporter genes are located within 5 kb from their putative target gene^87,88^. This is in stark contrast to gene regulation in mammalian cells, in which enhancers can be located megabases away from their target gene, but their activation potential is limited by TAD structure^89^.

In hermaphrodite nematodes, TADs are only present on the X chromosome due to the loading of the X-specific condensin I variant condensin I^DC^ on this chromosome^45,90^. Our study demonstrates that cleavage of condensin I/I^DC^ results in the loss of all TADs on the X chromosome. *In vitro,* condensins extrude DNA or chromatin loops^15,17^, and *in silico* modeling has shown that TAD structure formation requires dynamic loop extrusion that is limited by boundaries^20^. Our results, together with recent findings showing that the ATPase activity of DPY-27 is necessary for its binding to the X chromosome^91^, strongly suggest a model in which the SDC complex recruits condensin I^DC^ to *rex* sites and initiates loop extrusion, leading to the formation of TADs. Surprisingly, the cleavage of condensin I/I^DC^ reinforces a loop compartment formed by TAD boundaries, which are loading sites for condensin I^DC45^. These 35 loci create reciprocal loops upon cleavage of condensin I/I^DC^, spanning across genomic distances of 8-12 megabases. These loci coincide with *rex* sites, the loading sites for condensin I^DC^ ^92^, and show high enrichment for individual subunits of the SDC complex necessary for X-specific enrichment of condensin I^DC^ ^67^. A similar reinforcement of loops between TAD boundaries forming TAD corner peaks has been observed in mammalian cells upon knock-down of the cohesin destabilizer WAPL or protection from WAPL by acetylation of cohesin^STAG1^ ^93,94^. Mechanistically, long-range loop extrusion has been proposed to cause these horizontal and vertical dotted lines on HiC contact maps. However, in our experiments, condensin I/I^DC^ is released from chromatin, while in the studies mentioned above, loop extruders are stabilized onto chromatin. Therefore, long-range extrusion is unlikely to be the cause of the loop compartment formation observed in our study. To our knowledge, only one other study in mammalian cells describes the appearance of a loop compartment upon cohesin depletion, visualized by aligned dots on HiC maps^11^. In that study, loop anchors are a clique of strong, clustered enhancers known as “super-enhancers”, which form liquid-like condensates and are bound by the transcriptional co-activators BRD4 and MED1^95^. Interestingly, our data show that two of the SDC proteins form nuclear bodies, independently of condensin I/I^DC^ integrity, and are highly enriched at the TAD boundaries/loop anchors, which form the loop compartment upon condensin I/I^DC^ cleavage. Further experiments are required to decipher whether SDC bodies are indeed condensates.

Condensin I/I^DC^ cleavage leads to specific transcriptional upregulation of X-linked genes in nematodes, while only a few genes are deregulated on autosomes, with more genes significantly up- than down-regulated (366 vs 146 autosomal genes, respectively, Fig. 5a). Similarly, recent studies in budding and fission yeast have shown that condensin depletion or cleavage has little effect on transcription^23,96^, and these effects are believed to be mostly indirect due to chromosome missegregation during mitosis. However, in response to environmental stimuli such as starvation or quiescence, condensin has been observed to down-regulate gene expression, as in nematodes^97,98^. Regarding metazoans, it is less clear whether and how condensins play a role in gene regulation. In *Drosophila*, early reports suggest that the condensin I kleisin is necessary for *Fab-7* Polycomb responsive element repression^99^. More recent studies have identified the CAP-G subunit of condensin I as important for proper neuronal gene expression, yet genes were equally up- and down-regulated and the involvement of the entire condensin I complex remains unclear^100^. In our study, condensin II cleavage led to only a few misregulated genes without any chromosome specificity, with more genes being up- than down-regulated as for condensin I/I^DC^ cleavage (223 *vs* 65 genes, all chromosomes, Fig. 5a). These misregulated genes might result from an indirect effect of failed mitosis as they correlate strongly with genes misregulated upon cleavage of the cohesin^SCC-1^ complex which is solely expressed in dividing cells^32^ (Fig. 5c). Several studies have shown a role for condensin II in gene regulation in other experimental systems, yet it is less clear whether the complex is involved in repression or activation. The whole complex is involved in transvection by repressing gene expression *in trans*^101^ while the CAP-D3 subunit is necessary for activation of anti-microbial peptide expression^102^. Similarly, the murine CAP-G2 condensin II component is essential for erythroid cell differentiation and appears to repress gene expression^103^, but whether this is mediated by the condensin II holocomplex remains unclear.

In mammals^11,12^, studies have shown that lack of cohesins has little transcriptional effect on gene expression. Similarly, cleavage of the major *C. elegans* long-range loop extruder, condensin I, has little effect on autosomal genes. However, the absence of dosage compensation following condensin I^DC^ cleavage leads to the upregulation of genes located on the X chromosome. This highlights the importance of having a continuous presence of condensin I/I^DC^ on X chromatin to maintain X-linked gene regulation. Interestingly, the function of X chromosome TADs in *C. elegans* seems to be different from mammalian TADs, which limit the search space of enhancers^5,10^. Notably, in the worm, an equal number of cells expressed the X-linked gene *unc-3* under both control conditions and upon condensin I/I^DC^ cleavage (data not shown). Thus, removing TADs did not lead to ectopic activation of this gene by nearby enhancers situated in neighboring TADs. The imbalance in gene expression created by condensin I^DC^ cleavage between X chromosomes and autosomes is a major burden for the hermaphrodite animals and drastically reduces their lifespan. Strikingly, upon condensin I/I^DC^ cleavage, dosage compensation is maintained in the vicinity of loop anchors. These loci are recruitment sites for the SDC complex and have extremely high enrichment of SDC proteins as measured by ChIP-seq. They are, therefore, likely included in SDC bodies. Possibly, the function of loop extrusion by condensin I^DC^ could be to reel chromatin through the SDC bodies, which in turn would downregulate gene expression by an uncharacterized mechanism.

In conclusion, our study presents compelling evidence supporting the functional substitution of cohesin by condensin I in holocentric nematodes for interphase genome folding. Further characterization in other holocentric species will help determine if the use of condensin I is unique to *C. elegans* or if convergent evolution has led to similar mechanisms in other holocentric organisms^104^.

## Acknowledgments

We would like to thank Cihan Elcin and Michael Berger for technical help, Dr. Luca Giorgetti, Pr. Susan Gasser, Pr. Jeroen de Ridder, Dr. Amin Allahyar and Dr. Adriana Gonzalez for discussions and optimizations of the project, Dr. Pamela Nicholson and her team at the NGS platform of the University of Bern, the Meister laboratory for discussion, Dr. Kobus von Unen and the Pertz laboratory for help with HiLo microscopy. Some strains were provided by the CGC, which is funded by NIH Office of Research Infrastructure Programs (P40 OD010440). This work was funded by the Swiss National Science Foundation 31003A_176226/PP00P3_159320, the University of Bern, the SBFI ESKAS Program 2020.0321, and the Novartis Foundation for Medical/Biological Research.

## Methods

### Creation of TEV cleavage sites in individual kleisins and degron tagging of SDC-3

TEV cleavage sites were integrated into kleisin subunits of the different SMC complexes using CRISPR/Cas9 genome engineering^105^, using *unc-58* or *dpy-10* co-CRISPR as marker. Similarly, *sdc-3* was modified N-terminally by integrating the degron sequence in frame with the protein in the HW2079 strain (kindly provided by Dr. Helge Grosshans) expressing the TIR1 ubiquitin ligase under the control of the *eft-3* promoter. Template primers and guide RNAs are described in Supplementary Table S1.

### Phenotypic characterization

Animals synchronized at the first larval stage were grown at 22°C on fresh NGM plates seeded with OP50-1 bacteria. After 3 hours, the animals were heat-shocked at 34°C for 30 min and incubated at 22°C for the rest of the experiment, while they were imaged and their phenotype quantified.

### Western blot of TEV cleavage

Approximately 80,000 L1 synchronized worms were grown on 150 mm NGM plates and heat-shocked as described above. L3 stage worms were collected and washed three times with M9, and the worm pellet was frozen at −80°C. Pellets were defrosted and resuspended in 1 ml of 1xNPB buffer (20 mM Hepes pH 7.6, 20 mM KCl, 3 mM MgCl_2_, 0.5 M sucrose) supplemented with protease inhibitors, and 1 mM DTT. The worms were fragmented in a Balch homogenizer (Isobiotec) using 35 strokes with the 10 *μ*m ball and combined with two rinses of the homogenizer with 0.75 1xNPB. Fragmented worms were collected by centrifugation at 3200 g for 5 min at 4°C. An equal volume of 2xSDS-NaCl buffer (50 mM Tris, 2% SDS, 0.5 M NaCl) was added to the pellet, and samples were boiled for 5 min, followed by sonication in a Bioruptor machine (Diagenode) for 10 min, set on high, 30 s on, 30 s off. Insoluble material was removed from the lysate by centrifugation for 5 min at room temperature at 5000 g. For one batch of worms, 15,000 worms were treated +/− heat shock as above, and when collected they were directly lysed in 2x Laemmli buffer without Balch homogenization, before being boiled and sonicated as above. Samples were run on 4-12% ExpressPlus PAGE gels with Tris-MOPS-SDS running buffer (GenScript). Transfer was carried out in the cold room overnight in Tris-glycine buffer (250 mM Tris, 1.92 M Glycine, 10% ethanol) at 45 mA / 30 V onto nitrocellulose membrane. 3xFLAG-tagged kleisins from strains PMW1005(*dpy-26^cs^*), PMW1023 (*kle-2^cs^*), PMW1021 (*scc-1^cs^*) and PMW1025 (*coh-1^cs^*) were detected using the mouse monoclonal anti-flag M2 antibody (Sigma F1804). The heat-shock-inducible, myc-tagged TEV protease was detected with a mouse monoclonal anti-myc antibody (Sigma M4439). The proteins were detected using SuperSignal™ West Femto Maximum Sensitivity Substrate (ThermoFisher) and an Amersham Imager 600. The signals were quantified using ImageJ. For one blot where quantification was carried out for KLE-2 using a different exposure time, its signal intensity was normalized by the difference in the length of exposures. In all cases, band intensity was first normalized to the housekeeping gene tubulin (mouse anti-tubulin, Sigma T9026), and then in order to compare between blots, signals were normalized either to COH-1 present on the same blot or by normalizing heat shocked samples to non-heat shocked samples.

### Quantification of seam cell divisions

To evaluate the progression of seam cell division, strains with TEV sites in kleisins expressing a fluorescent seam cell marker (wls[scm::gfp] V) were treated as above, and the number of seam cells per animal was counted every 24 hours for the next three days.

### RNA-seq

For strains PMW366, PMW382, PMW775, PMW784, PMW828, and PMW844 synchronized L1 nematodes were seeded on NGM plates and incubated for 3 hours at 22°C. The animals were heat-shocked at 34°C for 30 min to activate the TEV protease and incubated again for 19 hours at 22°C. For strains PMW821, PMW823, PMW822, synchronized L1 nematodes were seeded on NGM plates with 1 mM auxin (prepared in ethanol). As a control, the strain PMW822 was seeded on NGM plates without auxin but with ethanol. L1 worms were incubated for 3 hours at 22°C, heat-shocked at 34°C for 30 min, and incubated for another 19 hours at 22°C. At that stage, most of the population reached the third larval stage. Animals were washed thrice with M9 buffer. RNA was isolated from the worms using Trizol, treated with DNase, and cleaned using RNeasy MinElute Cleanup Kit. Sequencing libraries were created using the Illumina Stranded mRNA preparation kit. Libraries were sequenced on an Illumina Novaseq 6000 device at the NGS platform of the University of Bern.

### RNA-seq data analysis

Between 37 and 60 million reads were generated from each library divided between two lanes of an Illumina NovaSeq 6000 machine. Adaptors were trimmed with Cutadapt v2.5, and reads were aligned to the WS275 version of the *C. elegans* protein-coding transcriptome using Salmon v1.5.2 in quant mode using –seqBias –gcBias and --numBootstraps 100 options. Differential expression analysis was carried out with DESeq2 v1.34. First, all genes with less than 10 reads in total among all samples were removed. The cleavage of each of the kleisin subunit was contrasted with the control strain expressing only the TEV protease, while accounting in the design formula for variation arising from sequencing lane, biological replicate, and sequencing date. Log2 fold change estimates were shrunken with the “apeglm” algorithm. An initial check showed that 35.1%, 51.5% and 49.6% of the significantly changing genes upon cleavage of DPY-26, KLE-2 and SCC-1, respectively, were genes that oscillate during larval development^106,107^. Oscillating genes could produce a large fold change in expression due to very small developmental asynchronies, therefore, we filtered out a combined list of 4522 genes that oscillate during larval development from the DEseq2 object before analysis. Basic processing of the RNAseq data and exploratory data analysis were carried out with custom scripts in bash and R, which can be found on github (https://github.com/CellFateNucOrg/SMC_RNAseq/tree/v0.2). The scripts used to generate the figures in this paper can be found at (https://github.com/CellFateNucOrg/MoushumiDas_paper/tree/v0.3.1). Significantly misregulated genes were filtered with an adjusted p-value of <0.05, and we considered either all statistically significant genes, or those that also had an absolute log2 fold change >0.5, as indicated in the figure legends. This threshold was chosen empirically to maximize the number of significant X-linked genes upon cleavage of DPY-26 while minimizing the number of autosomal genes that were considered significant.

RNAseq was performed in strains containing SDC-3 tagged with the auxin-inducible degron (AID) tag and TIR1 ubiquitin ligase with and without auxin, as described in Supplementary Table S5. For better comparison to previous samples, all strains contained the heat-shock inducible TEV protease and were heat-shocked at the L1 stage. Mapping and differential gene expression analysis was performed with Salmon and DESeq2 as described above. DESeq2 analysis was carried out by creating a dummy variable containing all combinations of *dpy-26^cs^*, *sdc-3^AID^*, TIR1, and auxin treatment that were present in the data and initially all strains were compared to the PMW366 control strain without auxin, taking into account sequencing lane, biological replicate and sequencing date as control variables. More complex comparisons were carried out in DESeq2 to remove the background effects of auxin and TIR1 by subtracting the coefficients of the different levels of the dummy variable. In the results, we focussed on the contrasts between PMW382 and PMW366 without auxin for *dpy-26^cs^* effects, and on the PMW822 strain with and without auxin for SDC-3 degron effects. The effect of the double mutant with both degradation of SDC-3 and cleavage of DPY-26 was estimated by subtracting the dummy variable coefficients for strain PMW823 with auxin from those of PMW821 with auxin in order to remove any non-specific effects of TIR1 and auxin in the absence of the degron.

### Lifespan assay

Animals were synchronized at the L1 stage and heat-shocked as above. When they reached the L4 stage, they were transferred to tight-fitting plates (BD Falcon Petri Dishes, 50×9mm) containing 50 mM FUdR and permanently shifted to 20°C. For degron-mediated degradation of SDC-3, *sdc-3^AID^*, half of the animals were transferred to FUdR plates with 1mM auxin diluted in DMSO, while the other half was transferred to FUdR plates with DMSO as a control. Plates were then loaded into air-cooled Epson V800 scanners and imaged with a frequency of two scans per hour using the lifespan machine setup^108^. The temperature of the scanner flatbed was continuously monitored (Thermoworks, Utah, US). Animals that exploded, burrowed, or escaped the imaging area were censored. For data processing, L4 stage animals were defined as day 0 of adulthood. The Kaplan-Meier product-limit procedure followed by the null hypothesis test employing the Mantel-Cox log-rank approach was applied to derive the survival function estimates. The survival data was processed in R using packages survminer (v0.3.1) and survival (v3.1-12).

Male lifespans are difficult to carry on plates as the animals tend to crawl off and die. To overcome this difficulty, mixed populations were bleached, hatched, and heat-shocked as above, and after two days at 20°C, 25 males were manually picked into 96 U-well microplates containing 100 µl S-basal with 5x concentrated *E. coli* per well, with 8 wells per strain as technical replicates. Health-span assays were performed using wMicroTracker ONE (MTK100) from InVivoBiosystems. Activity per well was measured once every 2 days until day 12 of adulthood and on a daily basis afterward. Each measurement included 3×30’ periods, with a 1-minute resolution. Motility rate of these periods was averaged between technical replicates for each strain. The experiment was repeated (biological replicates) and performed until reaching < 5% of starting motility rates in both conditions. Contaminated wells were excluded from the analysis. Data were analyzed using GraphPad® Prism 8 software.

### Construction of HiC libraries

3D chromatin conformation was acquired as in ref. 45, using synchronized animals 19 hours after heat shock for TEV induction as described above and with the following modifications. Animals were pelleted and stored at −80°C. For fixation, the frozen worm pellet was resuspended in 2% formaldehyde prepared in M9 buffer and incubated for 30 min in the rotator. Formaldehyde was then quenched with glycine at 125 mM (final), and incubated for 5 min. The worms were pelleted again, washed with M9, and about 100 ul of the packed animals were transferred to a 1.5 ml microcentrifuge tube, washed with 1 ml 1xPBS and protease inhibitors, washed again with 1 ml 1xPBS, snap frozen in liquid nitrogen before −80°C storage. Cross-linked frozen worms were ground using a SPEX6775 freezer/mill (SPEX Europe) at 5 cycles per second for 1 min with a total of 2 cycles. The ground worm powder was dissolved in 5 ml 1xPBS and immediately cross-linked with 500 μl TC buffer (100 mM NaCl, 1 mM EDTA, 0.5 mM EGTA, 50 mM Hepes pH 8.0, 22% formaldehyde), to a final concentration of 2% formaldehyde for 20 min at room temperature. Cross-linking was stopped by the addition of 289 μl of Stop Solution 1 (provided in the Arima HiC kit) and 5 min incubation at RT. Samples were pelleted at 2000 g for 15 min at RT and resuspended in 5 ml 1xPBS before 10 aliquots were distributed into 1.5 ml microcentrifuge tubes. The amount of DNA acquired from a single aliquot was estimated, and accordingly, the subsequent HiC experiment was done as per the manufacturer’s instructions. For library preparation, the modified protocol from Arima Genomics for the KAPA Hyper Prep Kit was used. The libraries were sequenced with Illumina sequencing to generate 100 bp paired-end reads. Sequence data is presented in Supplementary Table S3 and available from GEO (GSE199723).

### HiC data analysis

HiC Illumina data was processed using the HiCPro pipeline ^109^. A Singularity container containing version 3.0.0 of HiCPro was run with custom bash scripts with default parameters in the config file except for customization to the local cluster environment and the following parameters: BIN_SIZE = 2000, LIGATION_SITE = GATCGATC, GANTGATC, GANTANTC, GATCANTC for samples prepared with the Arima HiC kit, and using the ce11 genome downloaded from UCSC. The HiC-Pro results matrix was converted to cool format using the hicpro2higlass.sh tool from the Hic-Pro utils scripts. We then generated both balanced and unbalanced mcool files with our own custom resolutions (2000,4000,6000,8000,10000,20000,50000,100000,200000,500000 bp) with the cooler software (version 0.8.6) so they could be viewed in HiGlass. The custom bash scripts can be found in (https://github.com/CellFateNucOrg/hicpro/tree/v0.1). Additional data analysis was conducted with cooltools^110^, GENOVA^111^, and ad-hoc R scripts. For the conversion from eigenvector values to eigenvector values bins, the range of T/C or A/B eigenvector values (E1 or E2) determined for each genomic interval was split into 50 bins of equal genomic representation, and a bin value was assigned to each genomic interval, such as the lowest 2% of eigenvector values would get a bin value of 1 and the top 2% of the eigenvector values a bin value of 50. Spearman correlation of eigenvalues with expression data shown in Fig. 2C was carried out using all 2 kb regions with tpm>0. For 3-way contacts analysis, forward and reverse reads which did not aligned in the first round of “global” alignment of HiCPro were fused together *in vitro* and processed using the MC-HiC pipeline ^68^, using a 50 kb bin size and the bowtie2 mapper with parameters --very-sensitive -L 20 --score-min L,−0.6,−0.2 --end-to-end.

## Supplementary figure legends

**Figure S1.**
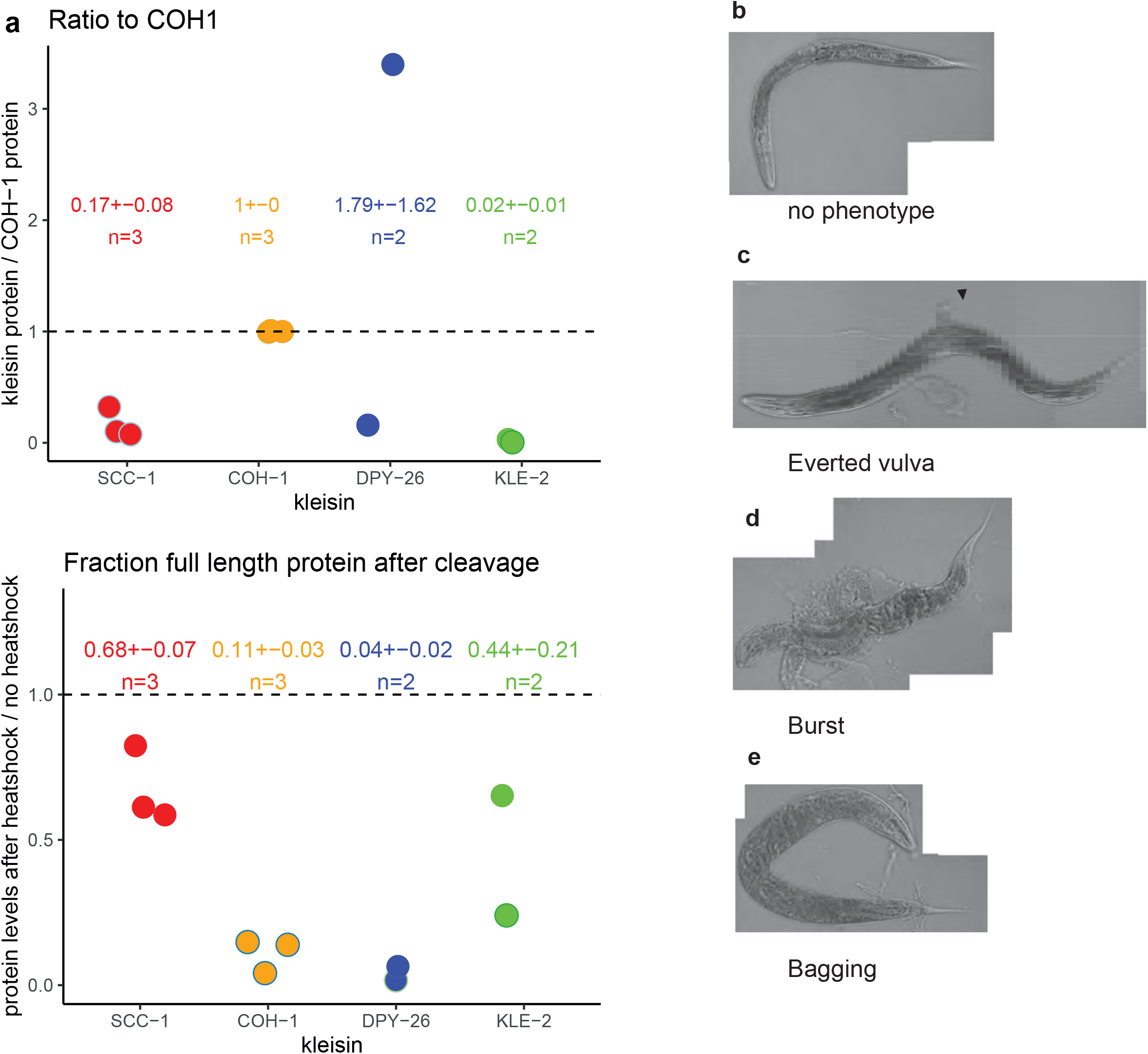
Kleisin cleavage quantification and phenotypes scored in animals 72 hours after kleisin inactivation. **a.** Quantification of kleisin cleavage by western blot. **Top panel:** Full-length FLAG-tagged kleisin protein levels (no heat shock, no TEV induction) were normalized to COH-1 levels on the same blot. Data are shown for 2 or 3 blots from biological repeats (n). **Bottom panel:** Ratio of FLAG-tagged kleisin protein levels 19 hours after TEV induction normalized to protein levels in age-matched controls without TEV induction. Mean +/− standard deviation of protein levels for each ratio are shown, as well as the number of biological replicates (n). **b.-e.** Scored phenotypes of animals 72 hours after kleisin cleavage induction. Pictures were taken with a 20x objective, 72 hours after SCC-1 cleavage. Wild-type animal (**b**). Everted vulva (marked with a black arrow, **c**). Animals burst through the vulva (**d**). “Bagging” or Egg-laying defective (Egl) animals in which a non-functional vulva leads to the retention of embryos inside the mother’s body (**e**).

**Figure S2.**
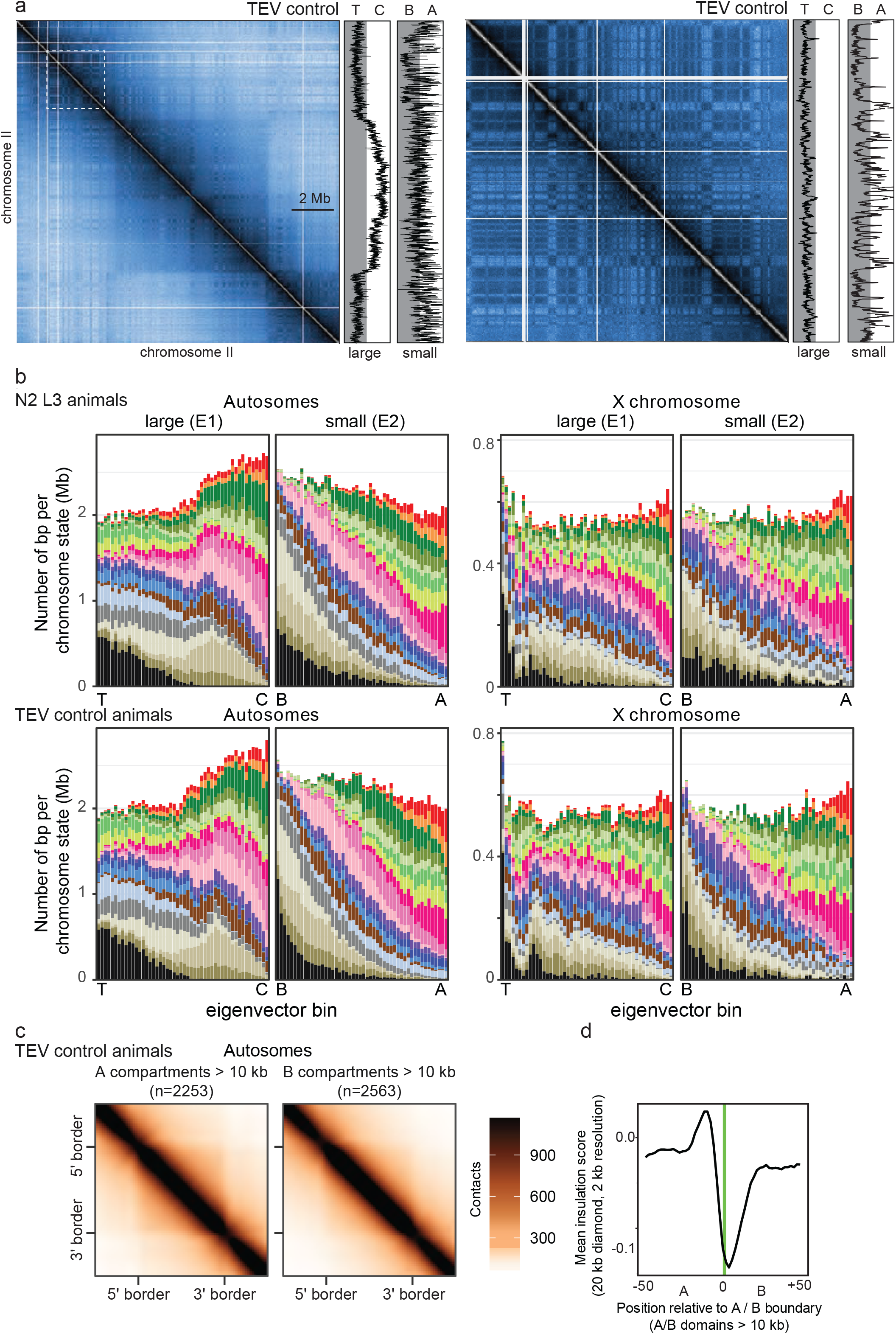
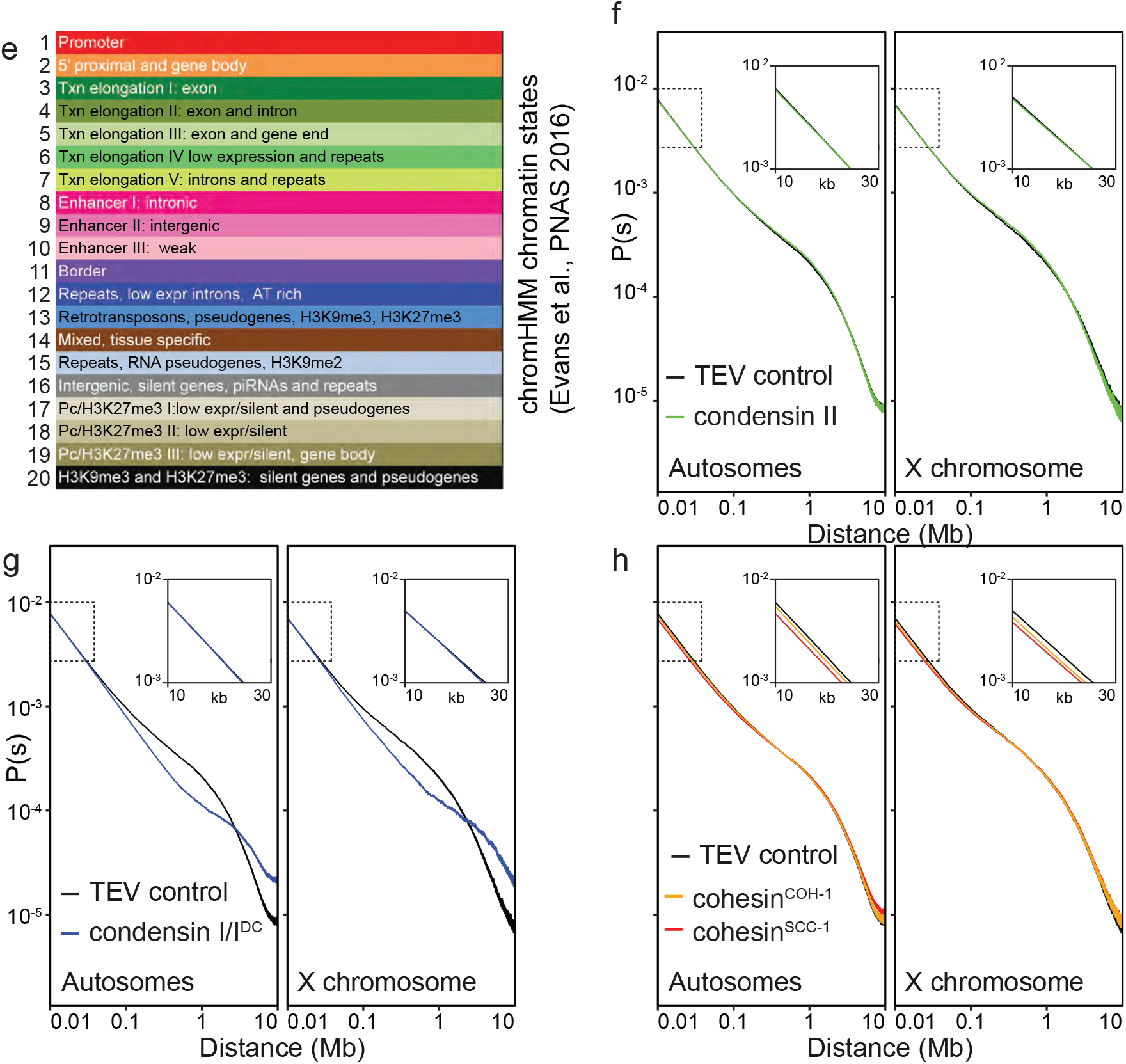
HiC captures compartmentation at different scales. **a.** HiC contact frequency map of chromosome II. Left: whole chromosome map and corresponding first (large) and second (small) eigenvectors determined on the genome-wide *trans* contact-frequency maps (right side). Same as Fig. 2a. Right: Detail of chromosome-wide map (dashed square on the left chromosome-wide contact map), with corresponding eigenvectors. Notice the correlation between the cross-hatched pattern and the values of the second (small) eigenvector. **b.** Chromatin state composition of eigenvector bins in wild-type N2 animals and TEV control animals (same as Fig. 2d for comparison between X and autosomes). The chromatin state composition of each chromosome interval as classified by ^58^, panel e, was correlated with the eigenvector value of each 2 kb interval for autosomes and the X chromosome. **c.** A and B compartments defined by the second *trans* eigenvector form insulated domains. Scaled average HiC contact map of A and B domains larger than 10 kb. **d.** Average insulation score from A to B compartments measured using a diamond of 20 kb at a resolution of 2 kb. The last A-type bin of the transition is marked in green. **e.** chromHMM states as classified by ^58^. **f.** Contact probability decay plots for autosomes and the X chromosome in TEV control animals and after kleisin cleavage of condensin II. **g.** Same as in F for TEV control animals and after kleisin cleavage of condensin I/I^DC^. **h.** Same as in F for TEV control animals and after kleisin cleavage of cohesin^SCC-1^ or cohesin^COH-1^.

**Figure S3.**
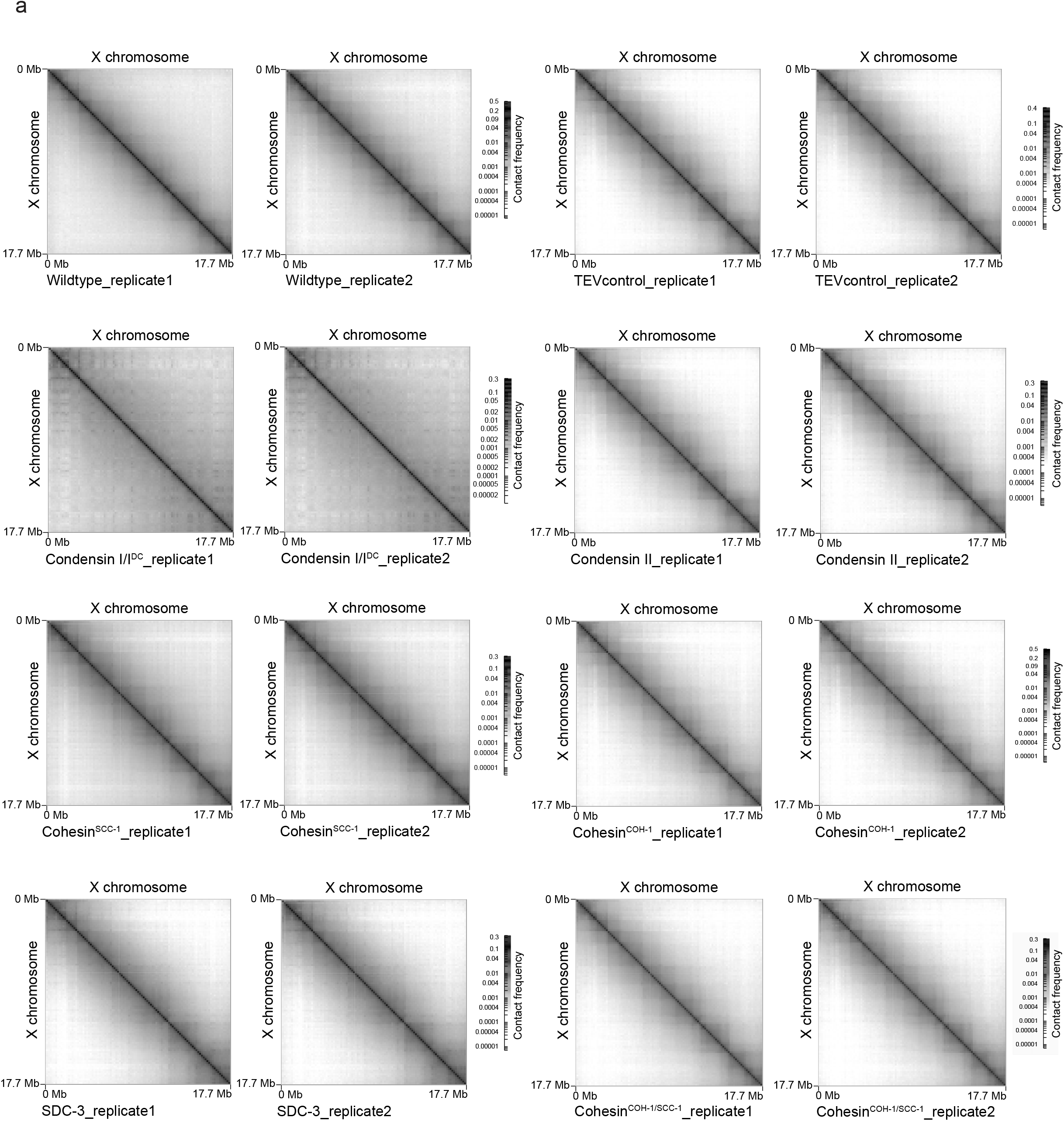
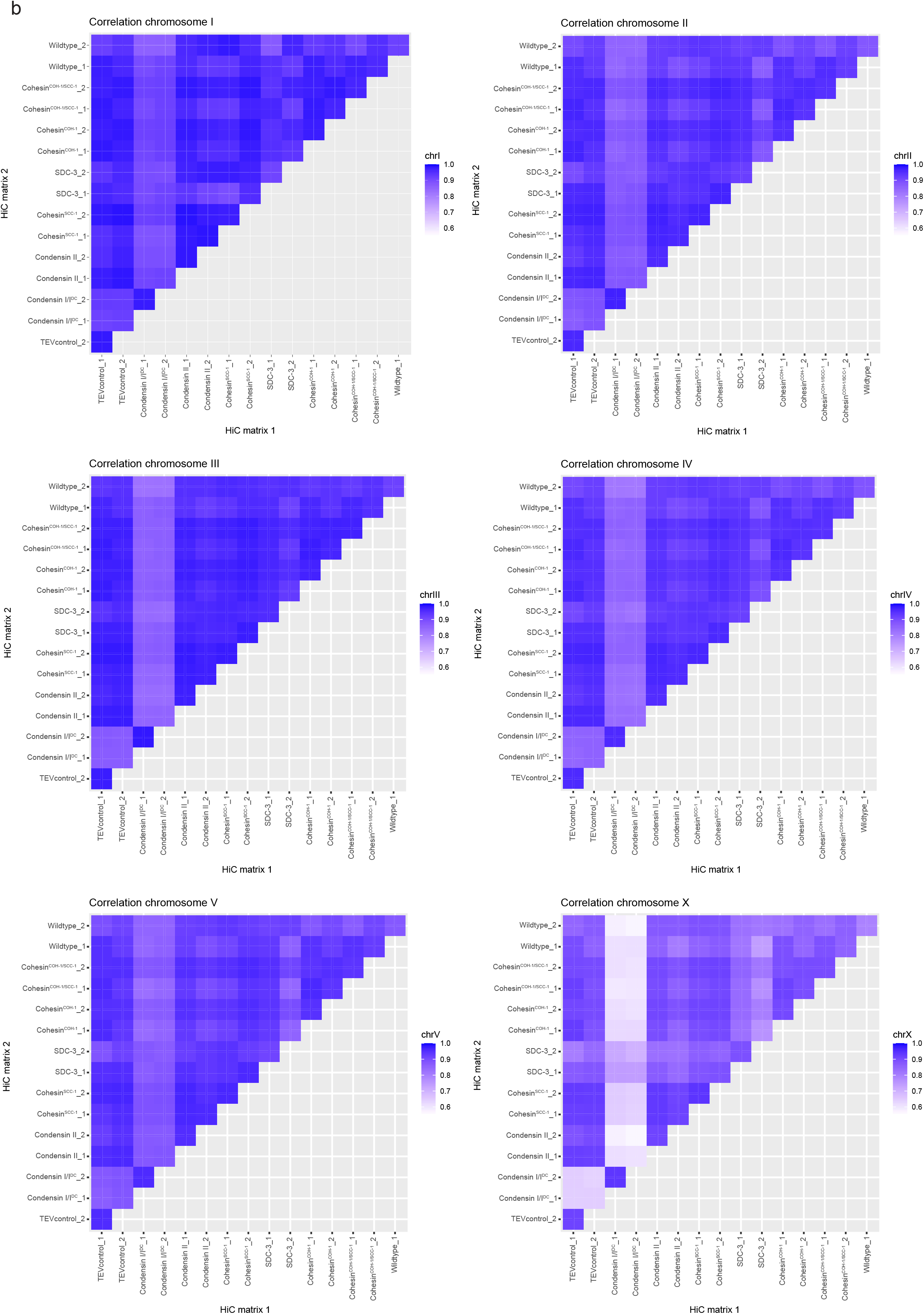
Comparison of the biological replicates from HiC experiments. **a.** HiC maps of the X chromosome from individual repeats in L3 animals. Libraries were processed using HiC-Pro^109^ and visualized using HiGlass^112^. The contact matrices of the X chromosome are shown for both biological replicates for each condition (wild-type animals, TEV control, and upon cleavage of condensin I/I^DC^, condensin II, cohesin^SCC-1^, cohesin^COH-1^, cohesin^COH-1/SCC-1^ and degron-mediated knock-down of SDC-3). **b.** Heat-maps quantifying HiC reproducibility for the different chromosomes calculated using HiCrep^113,114^.

**Figure S4.**
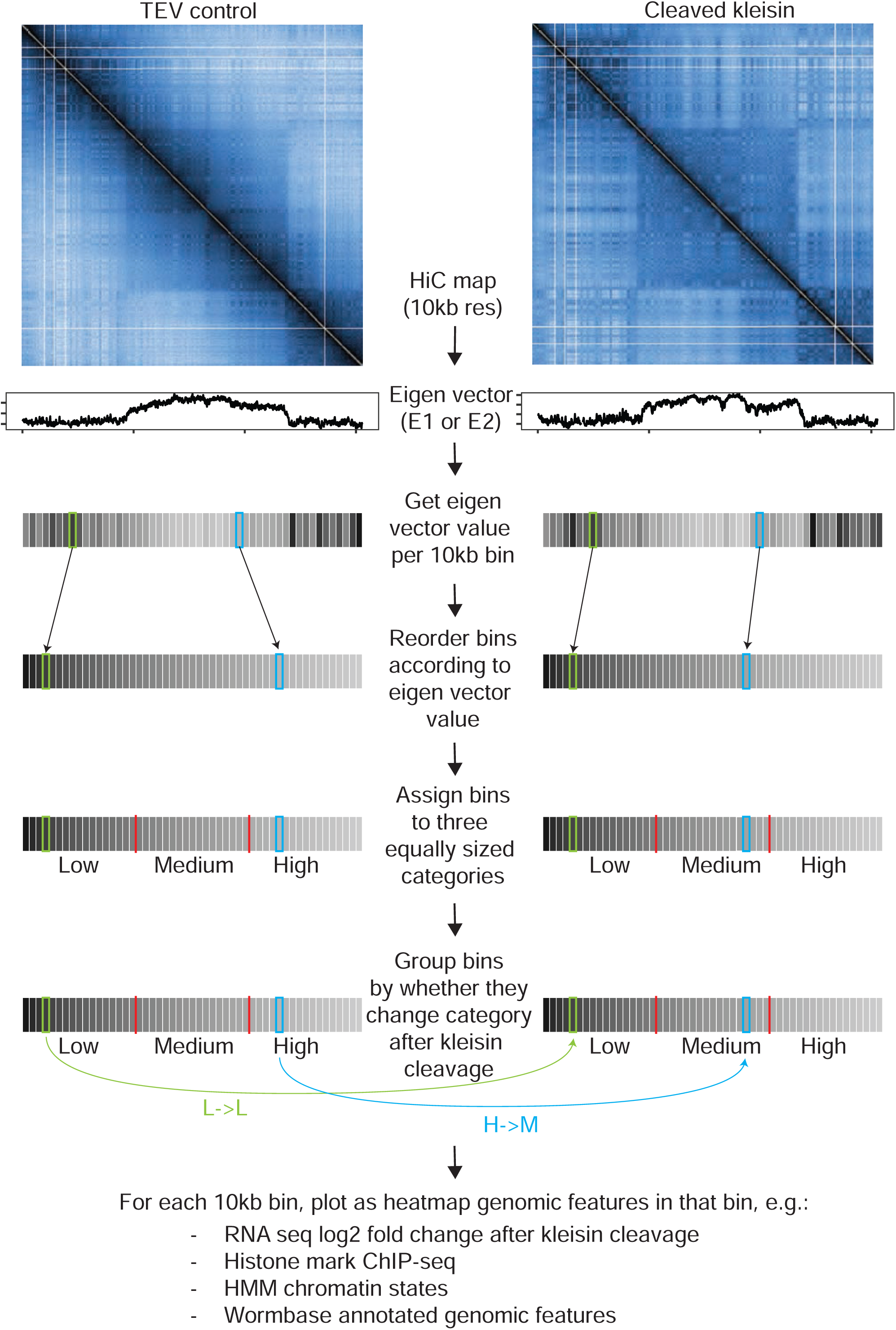
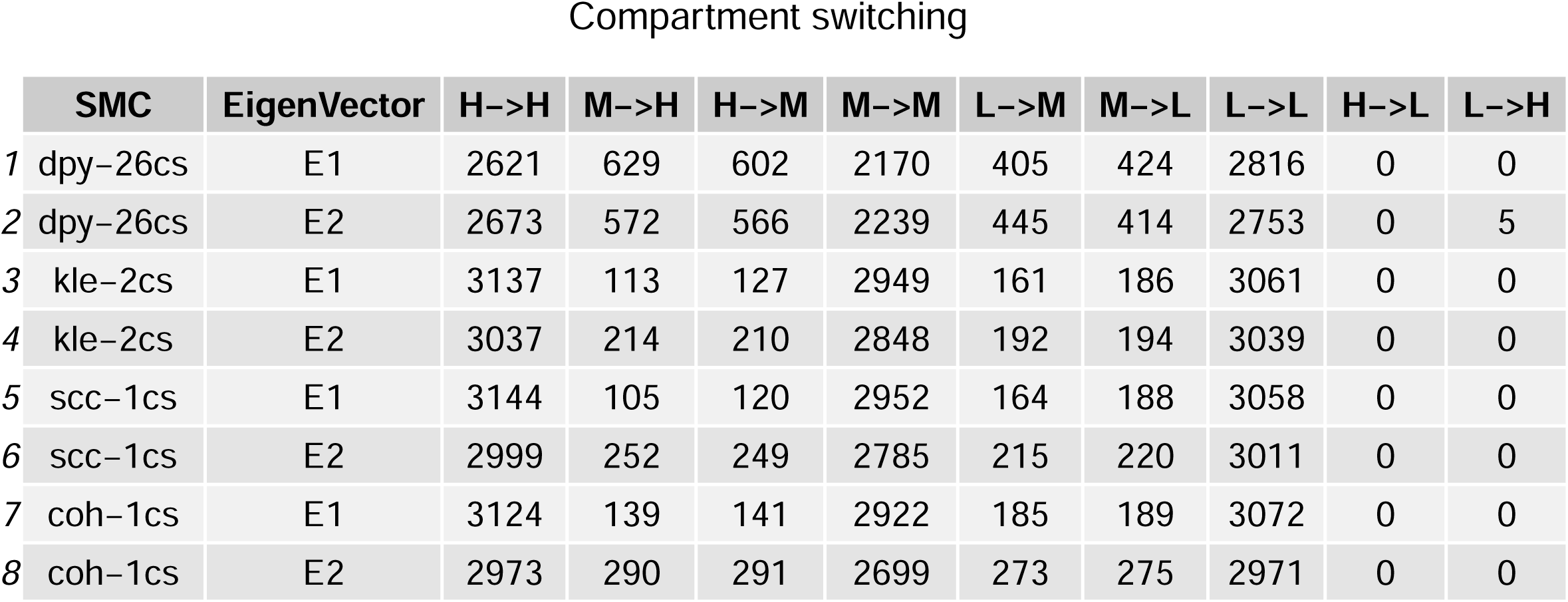
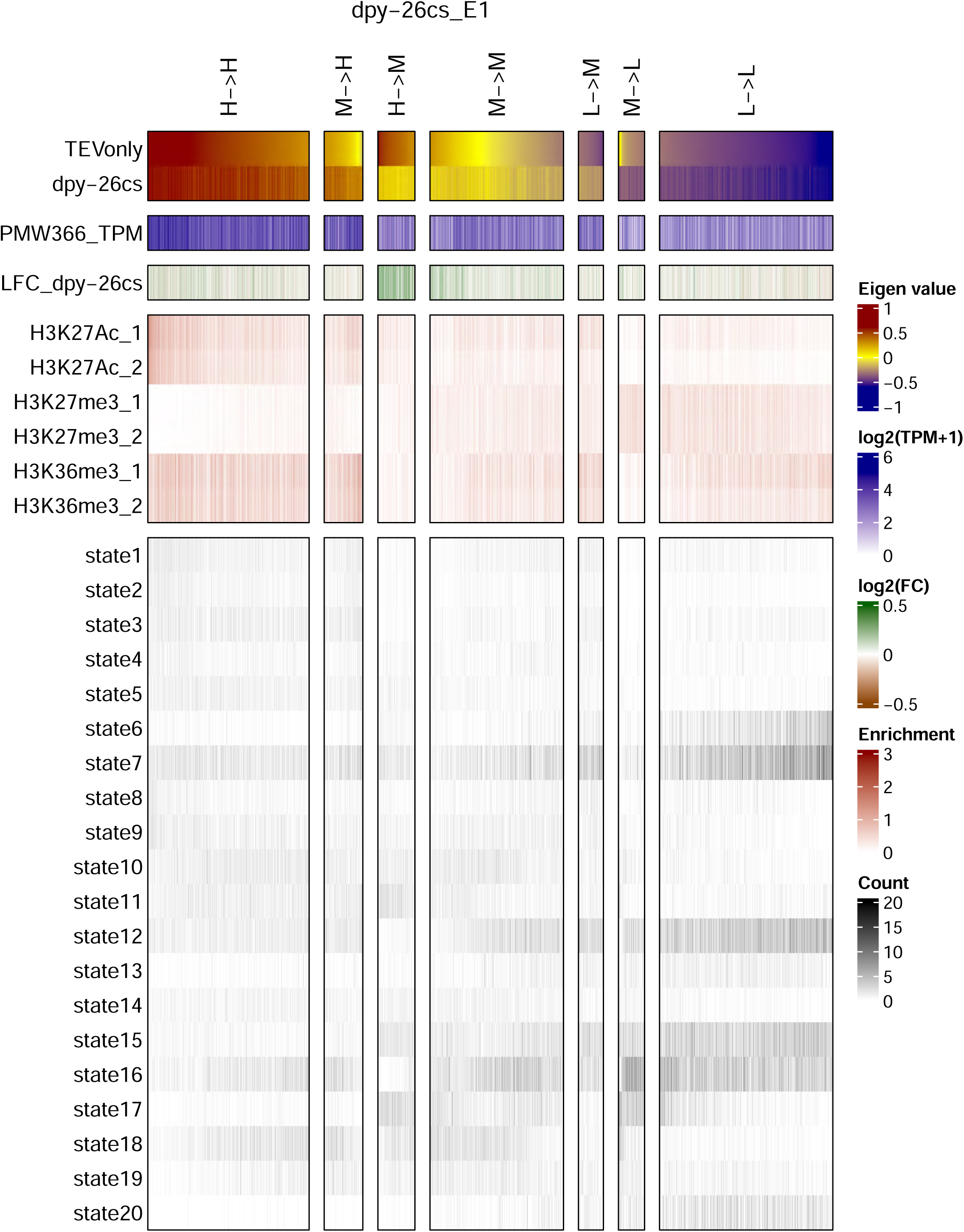
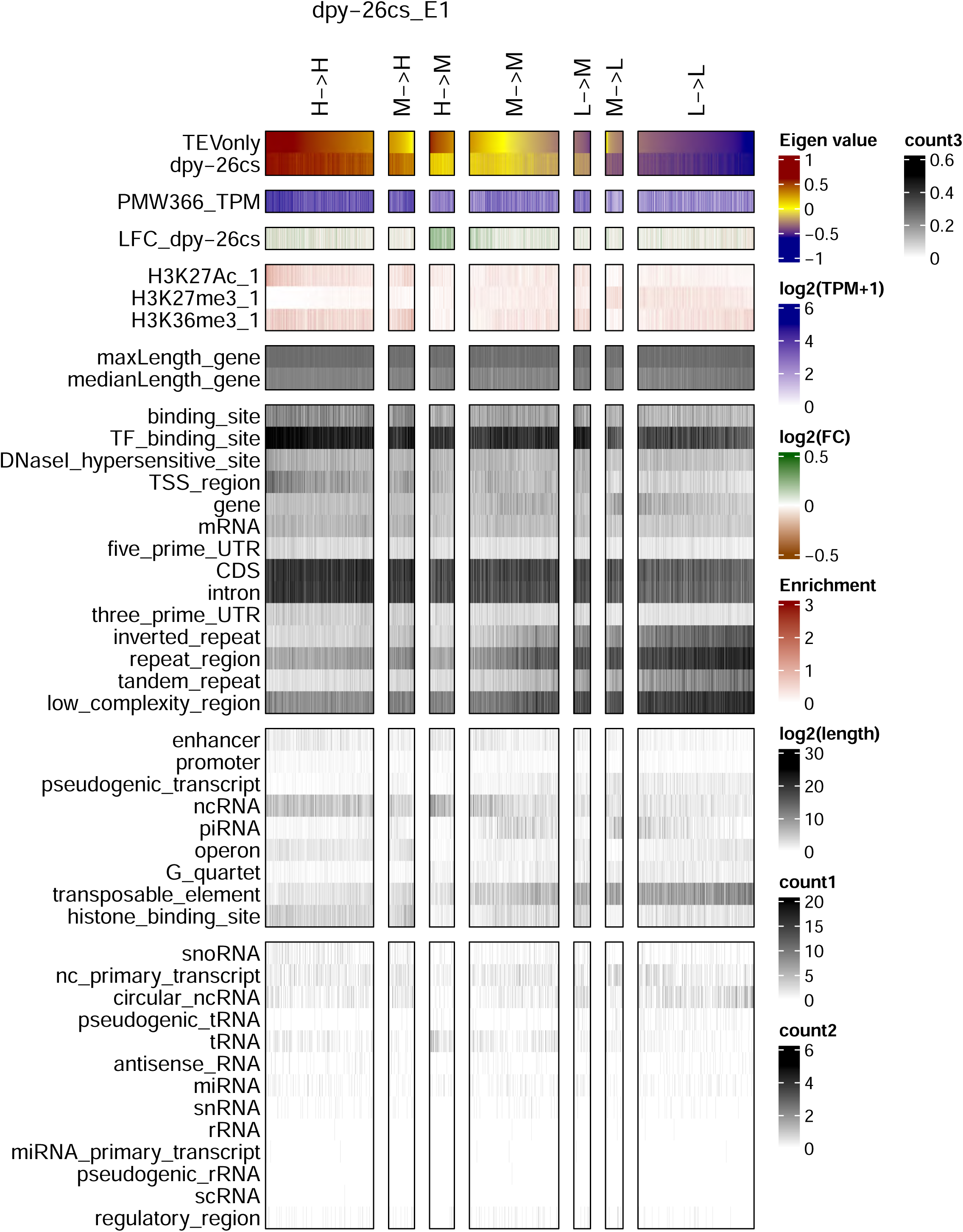
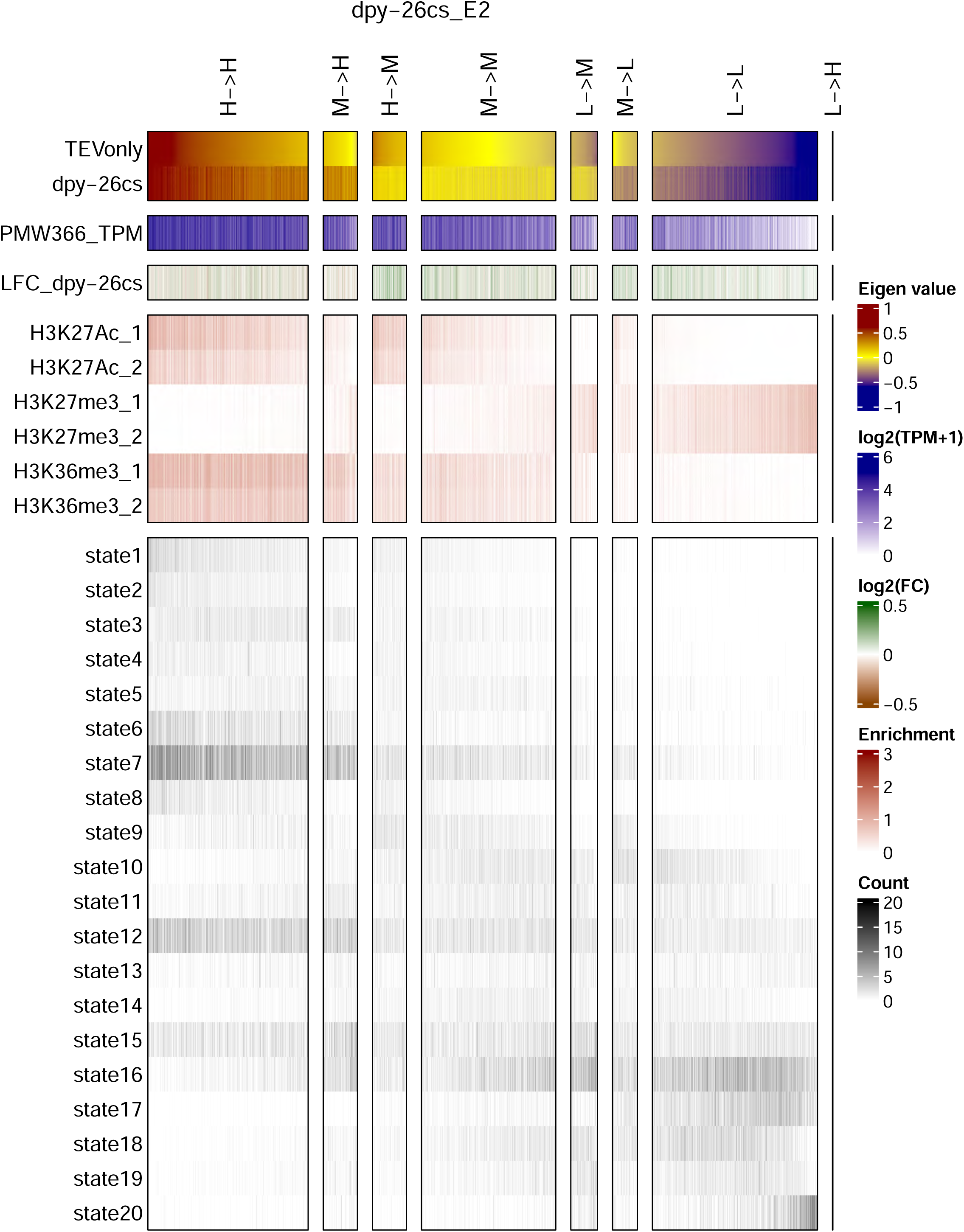
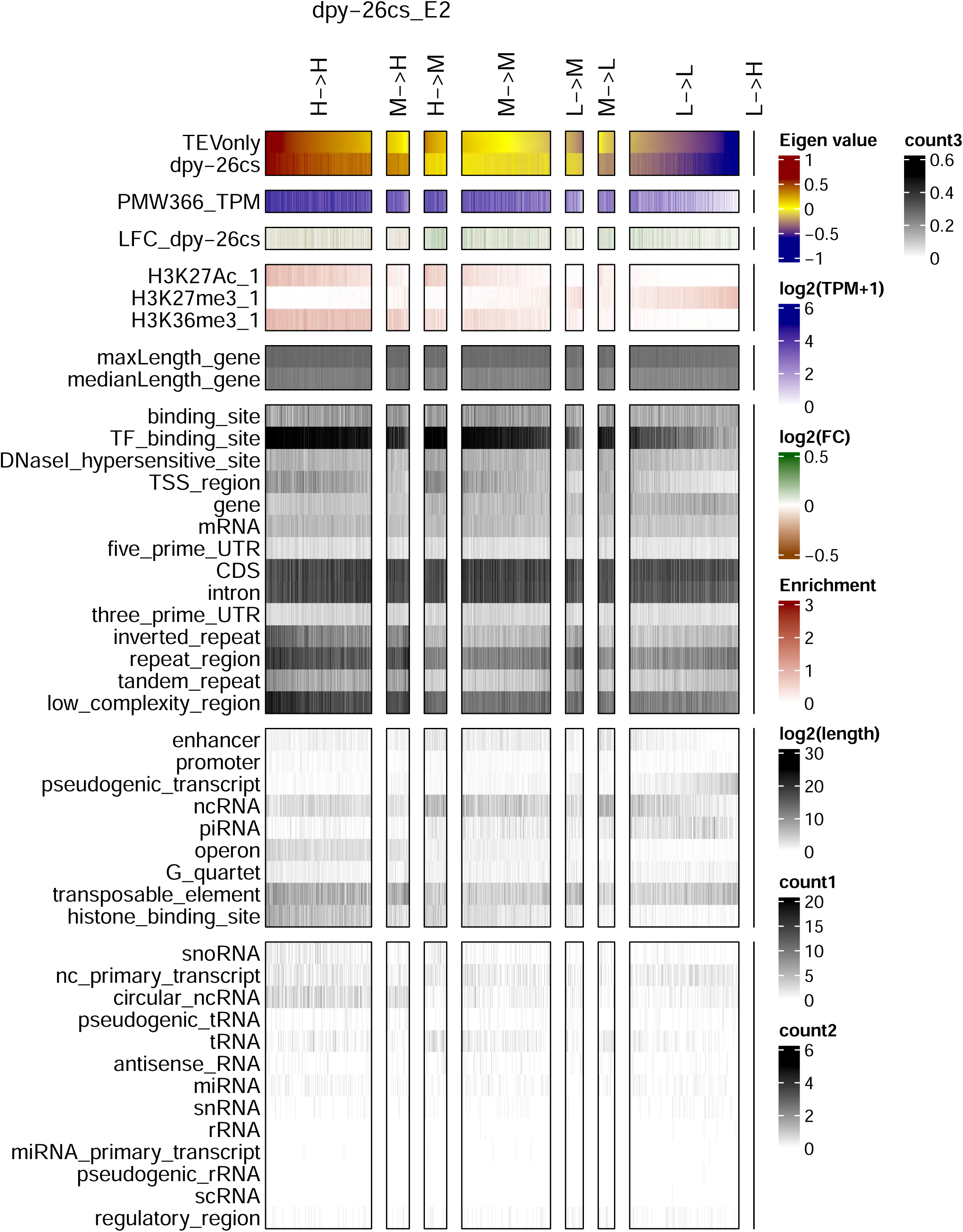
Epigenomic and genomic characteristics of HiC compartments upon kleisin cleavage. First page: diagram explaining the classification of eigenvector bins shown on subsequent pages. The first two eigenvectors (E1 and E2) of the 10 kb HiC matrix for each kleisin cleavage and the TEVonly control were divided into 3 quantiles, designated high (H), medium (M), and low (L). Each 10 kb genomic bin was scored according to which tercile it was in the TEV control experiment HiC sample and after cleavage of the kleisin subunit *dpy-26^cs^*(condensin I/I^DC^), *kle-2^cs^* (condensin II), *scc-1^cs^*(cohesin^SCC-1^) or *coh-1^cs^* (cohesin^COH-1^). The number of genomic bins that either remained in the same compartment (H->H, M->M or L->L), went from a higher to the lower compartment (H->M, M->L, H->L) or lower to higher (M->H, L->M, L->H) compartment upon kleisin cleavage are shown in the table on the second page. Subsequent pages show heatmaps with the bins from the condensinI/I^DC^ (*dpy-26^cs^*) eigenvectors clustered by type and sorted by the descending value of their eigenvector in the control sample (TEV control), with additional genomic/chromatin features scored in these bins. The first two heatmap pages contain heatmaps for the bins created from the E1 eigenvector, and the next two pages contain heatmaps for the bins created from the E2 eigenvector, as indicated at the top of the pages. **Top panel (red-yellow-blue):** Eigenvector values for the TEV control and kleisin cleavage datasets. **Second panel (white-blue):** Mean basal gene expression (transcripts per million) in the control sample (PMW366_TPM). **Third panel (red-white-green):** log2 fold change upon kleisin cleavage (LFC_). **Fourth panel (white-red):** Average ChIP seq signal per bin for histone marks: H3K27Ac (GSM624432, GSM624433), H3K27me3 (GSM562734, GSM562735) and H3K36me3 (GSM562736, GSM562737). Note that on heatmaps 2 & 4 only the first replicate is shown to save space. **Fifth panel on heatmaps 1 & 3 (white-black):** Count of the number of occurrences of the different HMM chromatin states defined by ^58^ in each bin. **Fifth panel on heatmaps 2 & 4 (white-black):** log2 of the maximum and median gene length in base pairs in each bin. **Sixth, seventh and eighth panels on heatmaps 2 & 4 (white-black):** Counts per bin of the occurrence of different genomic features extracted from the Wormbase WS280 genome annotation GFF3 file. The features were divided into three separate panels to account for the different abundance of the features.

**Figure S5.**
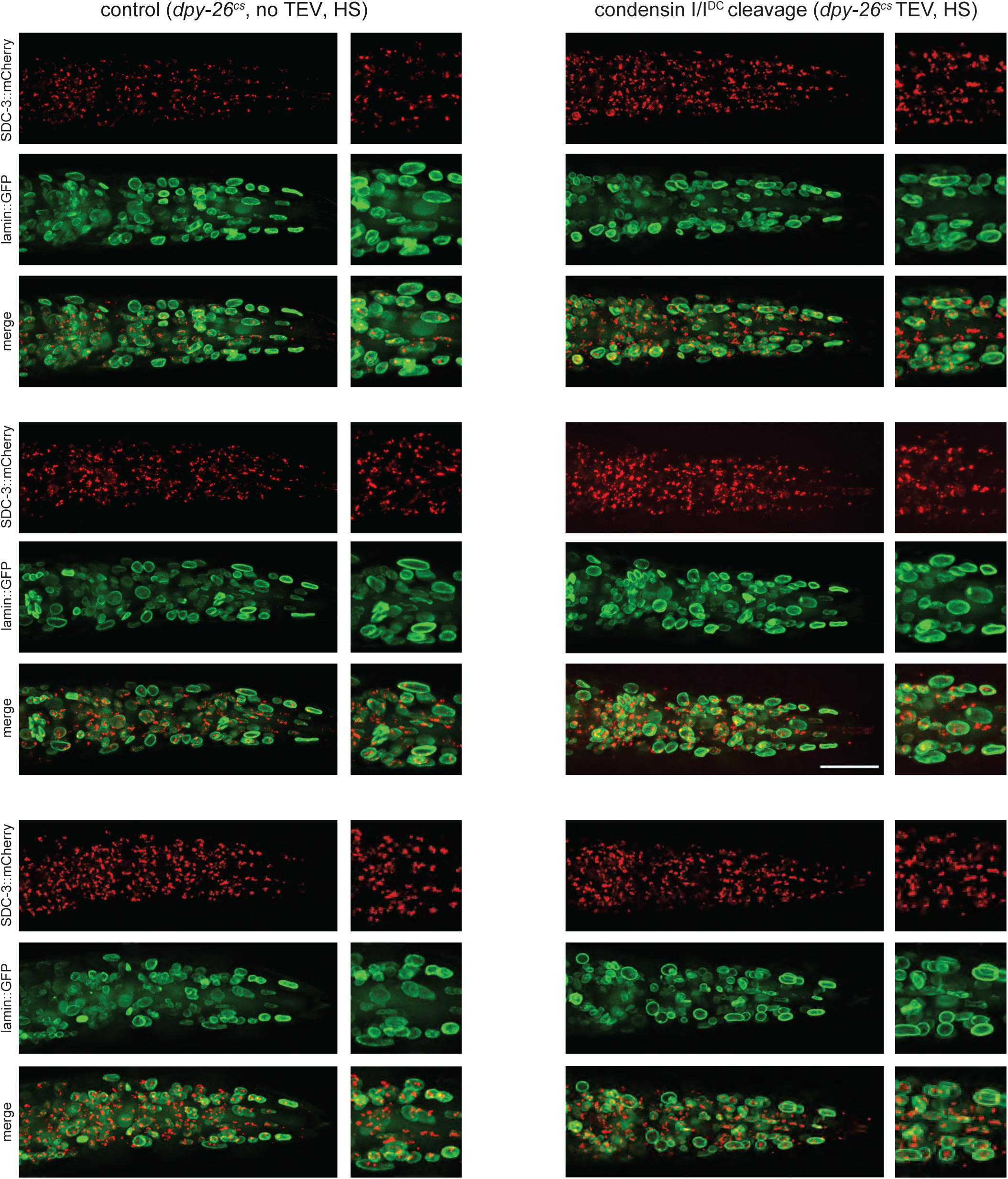
Absence of modification of the SDC-3::mCherry nuclear pattern following condensin I/I^DC^ cleavage. Fluorescence pattern in worms expressing the nuclear periphery marker GFP lamin (green) and SDC-3::mCherry (red) 19 hours post TEV protease induction by heat shock. Right column: control animals carrying the TEV^cs^ insertion in the *dpy-26* open reading frame but not the TEV expression construct. Left column: animals carrying both the TEV^cs^ in *dpy-26* and the TEV heat shock dependent expression construct. The SDC-3 fluorescence signal is indistinguishable between the two strains (assessed blindly). Synchronized L1 worms were kept at 22°C for 3 hours before a 30 minute heat shock at 34°C. Imaging was carried out 19 hours after heat shock. Scale bar: 20 μm.

**Figure S6.**
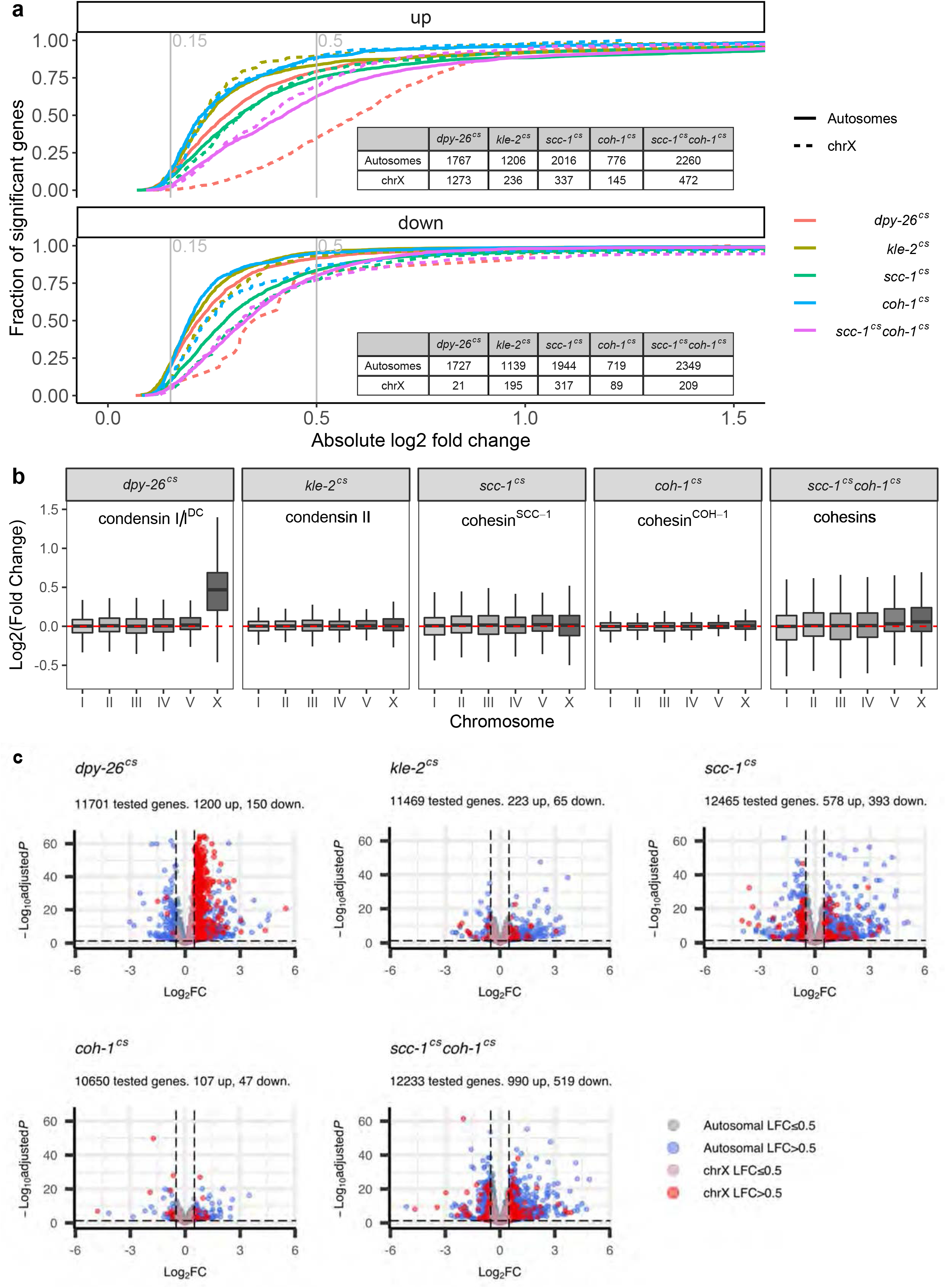
Differential gene expression after cleavage of kleisin subunits of different SMC complexes. **a.** Empirical cumulative distribution of the log2 fold change of significantly changed genes (AdjP<0.05, |LFC|>0) after cleavage of condensin I/I^DC^ (*dpy-26^cs^*), condensin II (*kle-2^cs^*), cohesin^SCC-1^ (*scc-1^cs^*), cohesin^COH-1^ (*coh-1^cs^*) or cohesin^SCC-1/COH-1^ (*scc-1^cs^coh-1^cs^*). Top panel shows upregulated genes, bottom panel shows downregulated genes. Vertical gray lines mark 0.15 and 0.5 log2 fold change which is the range that includes the log2 fold change for most of the significantly changing genes from all samples except X-linked genes up-regulated upon cleavage of condensin I/I^DC^ (*dpy-26^cs^*), dashed red line in the top panel. The number of significantly up and down-regulated genes for each sample and each type of chromosome is shown in the inset tables. **b.** Log2 fold change for 13734 genes with at least 10 read counts in total. Horizontal red line marks no change in gene expression. **c.** Volcano plots for all genes filtered by DESeq2 as expressed (“tested”, AdjP is not NA). X-linked genes are shown in red and dull red and autosomal genes are shown in blue and dull blue (|LFC|>0.5 and |LFC|≤0.5, respectively). The horizontal dashed line marks an AdjP value threshold of 0.05, and the vertical dashed lines mark LFC thresholds of −0.5 and 0.5. The number of statistically significant genes (AdjP<0.05 and |LFC|>0.5) in each dataset is shown in the plot sub-heading.

**Figure S7.**
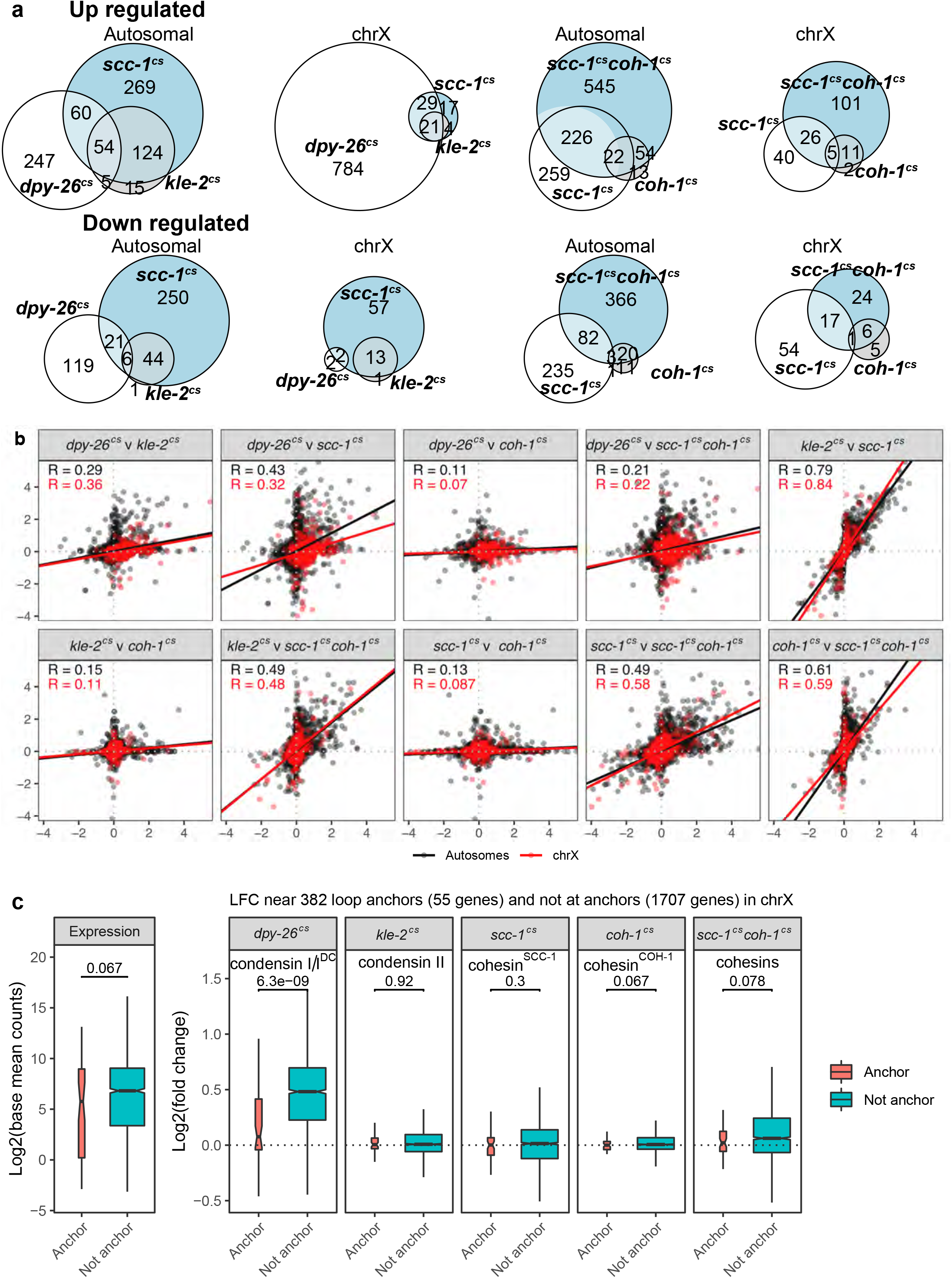
Comparison of differential gene expression changes between different datasets. **a.** Overlap of significantly changing genes (AdjP<0.05, |LFC|>0.5) after cleavage of different SMC complexes. First and second columns show overlap for condensin I/I^DC^ (*dpy-26^cs^*), condensin II (*kle-2^cs^*) and cohesin^SCC-1^ (*scc-1^cs^*) cleavage. Third and fourth columns show overlap for cohesin inactivation by targeting either single kleisins (*scc-1^cs^* or *coh-1^cs^*) or both (*scc-1^cs^coh-1^cs^*). **b.** Pairwise correlation of the log2 fold change for 13734 genes with at least 10 read counts in total. X-linked genes (1783) are shown in red and autosomal genes (11951) in black. The lines represent the best fit linear regressions and the Pierson correlation coefficient is shown in each panel. **c.** Comparison of 55 genes that overlap 10 kb regions around 35 loop anchors on the X chromosome (Fig. 3a, “Anchor”) defined using the second *cis* eigenvector of the HiC matrix upon condensin I/I^DC^ cleavage, with 1707 genes found on the X chromosome that did not overlap loop anchor regions (“Not anchor”). Leftmost panel shows DESeq2 base mean expression value, and the other panels show the difference in log2 fold change of expression for each dataset in these two sets of genes. The p-values from the Wilcoxon rank sum test comparing anchor to non-anchor regions for each dataset are shown in each panel.

**Figure S8.**
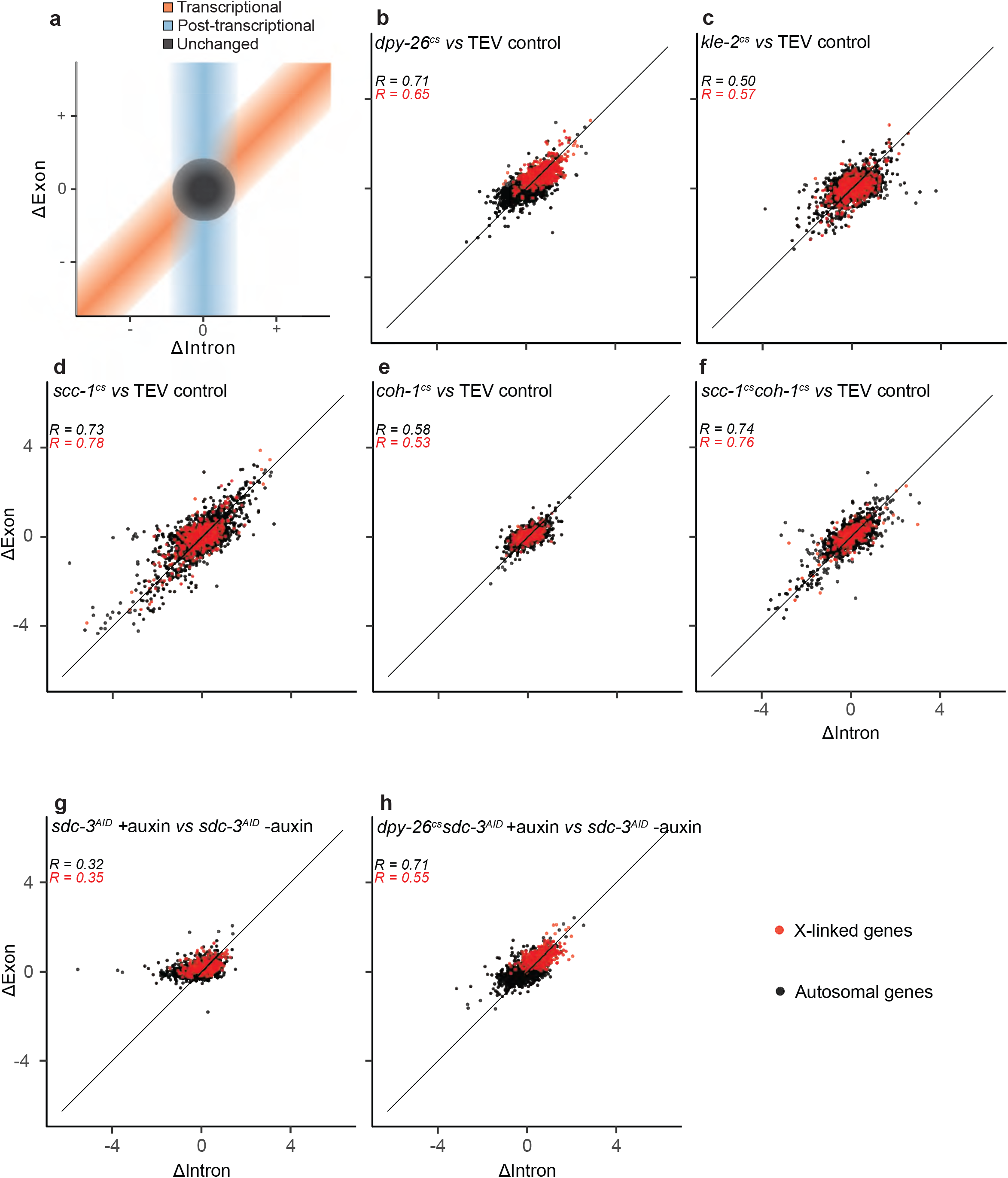
Exon-Intron Split Analysis (EISA) of transcriptional changes upon kleisin cleavage. **a.** Schematic explanation of the EISA. ΔIntron and ΔExon refer respectively to the intronic and exonic expression changes observed in the experiment. **b-f.** Comparisons of ΔIntron and ΔExon upon cleavage of the indicated kleisins. R designates the Pearson correlation coefficient of ΔIntron and ΔExon for autosomal genes (black) and genes on the X chromosome (red). **g.** Comparison as in b upon auxin-mediated knock-down of the condensin I^DC^ loader SDC-3 compared to control conditions without auxin. **h.** Comparison as in b between condensin I/I^DC^ cleavage with SDC-3 auxin-mediated degradation compared to control conditions without auxin. Key as in b-f.

**Figure S9.**
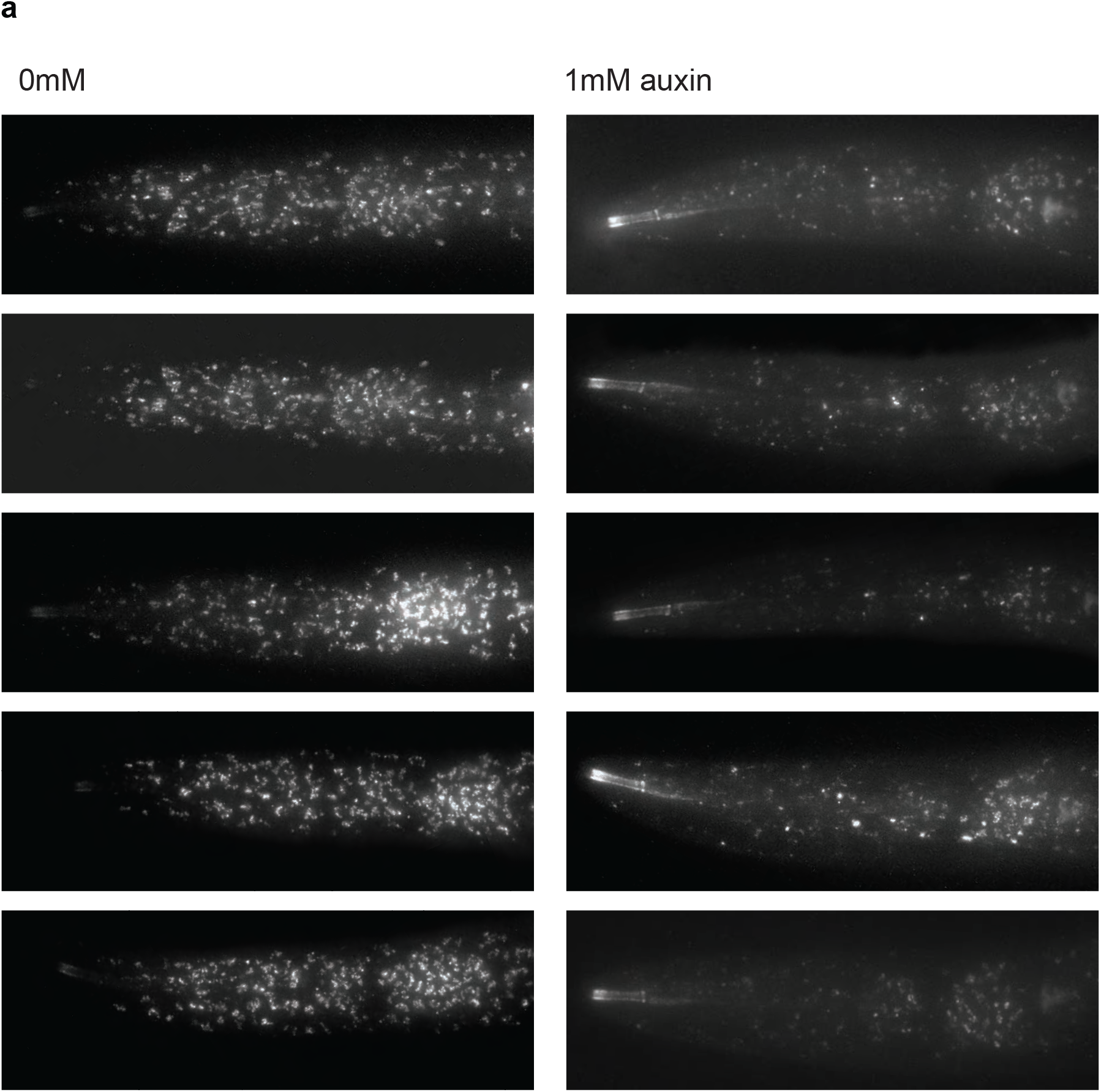
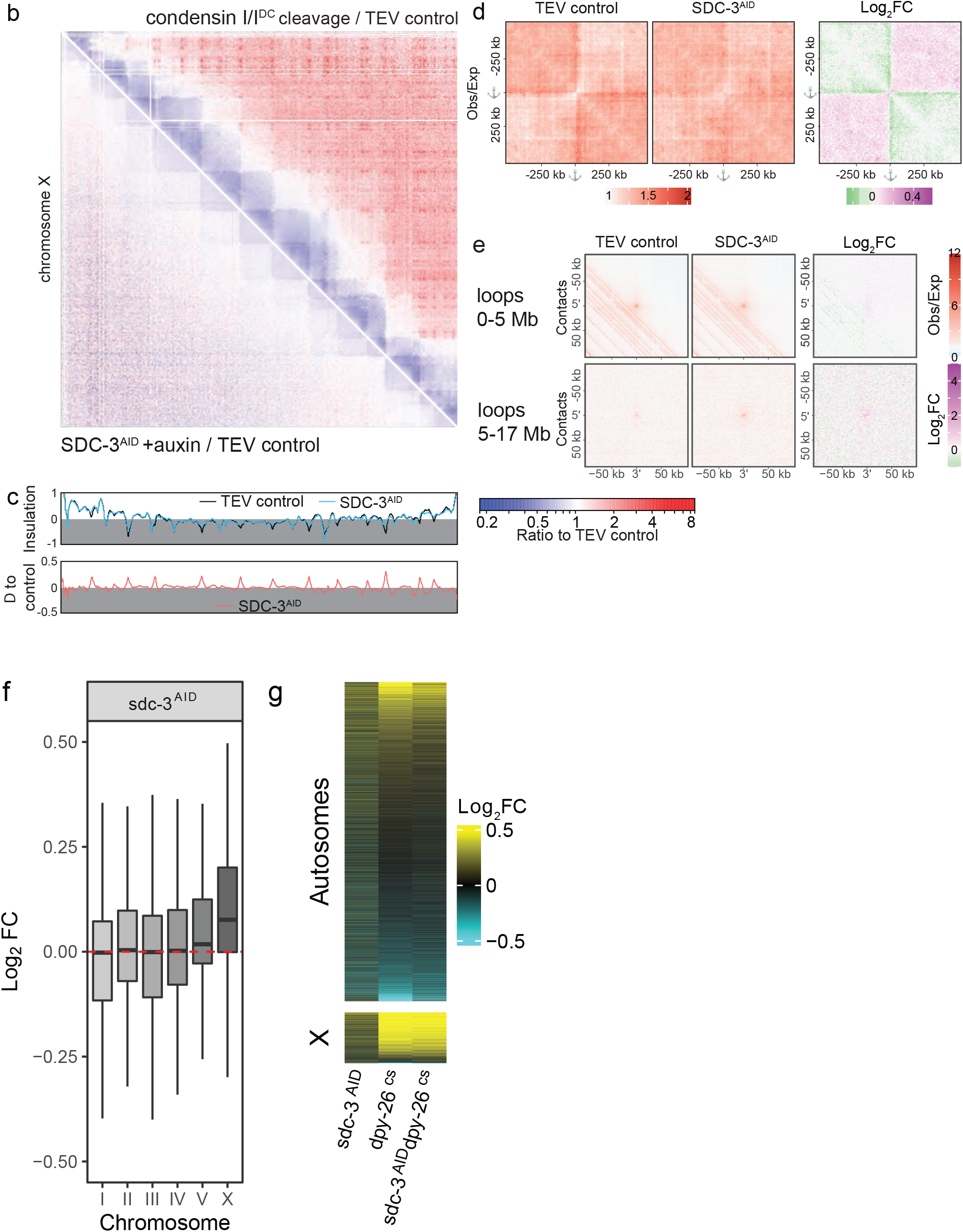
Effects of SDC-3 degradation on X chromosome structure and transcription. **a.** Auxin treatment leads to partial SDC-3 degradation. Synchronized first larval stage animals expressing degron-tagged SDC-3::mCherry were grown on medium without (left) or with 1mM auxin (right) and imaged after 18.5 hours at 25°C. **b.** HiC contact frequency ratio maps to TEV control after condensin I/I^DC^ cleavage (top right) and SDC-3 degron knock-down (bottom left) for the X chromosome. Pooled data from two biological replicates per condition. **c.** (top) Insulation score of the X chromosome for TEV control (black) and SDC-3 degron (blue). (bottom) Difference between SDC-3 knock-down (SDC-3^AID^) and control. **d.** Average HiC contact maps centered on the anchors defined in Fig. 4a, highlighting decreased TAD organization upon SDC-3 knock-down. **e.** Average HiC contact maps for all possible loops between pairwise combinations of anchors as in Fig. 4e. Loops are split by size between 0-5 Mb and >5 Mb. **f.** Log2 fold change of genes by chromosome upon SDC-3 knock-down (*sdc-3^AID^*), highlighting a modest, yet significant upregulation of X-linked genes. **g.** Hierarchical clustering of the LFC of 12465 genes filtered by DESeq2 as expressed in at least one dataset after knockdown of SDC-3 (*sdc-3^AID^*), cleavage of condensin I/I^DC^(*dpy-26^cs^*) or both (*sdc-3^AID^dpy-26^cs^*). Top panel: 10792 autosomal genes. Bottom panel: 1673 X-linked genes. Data for *dpy-26^cs^* is the same as in Fig. 5, included here for comparison.

**Figure S10.**
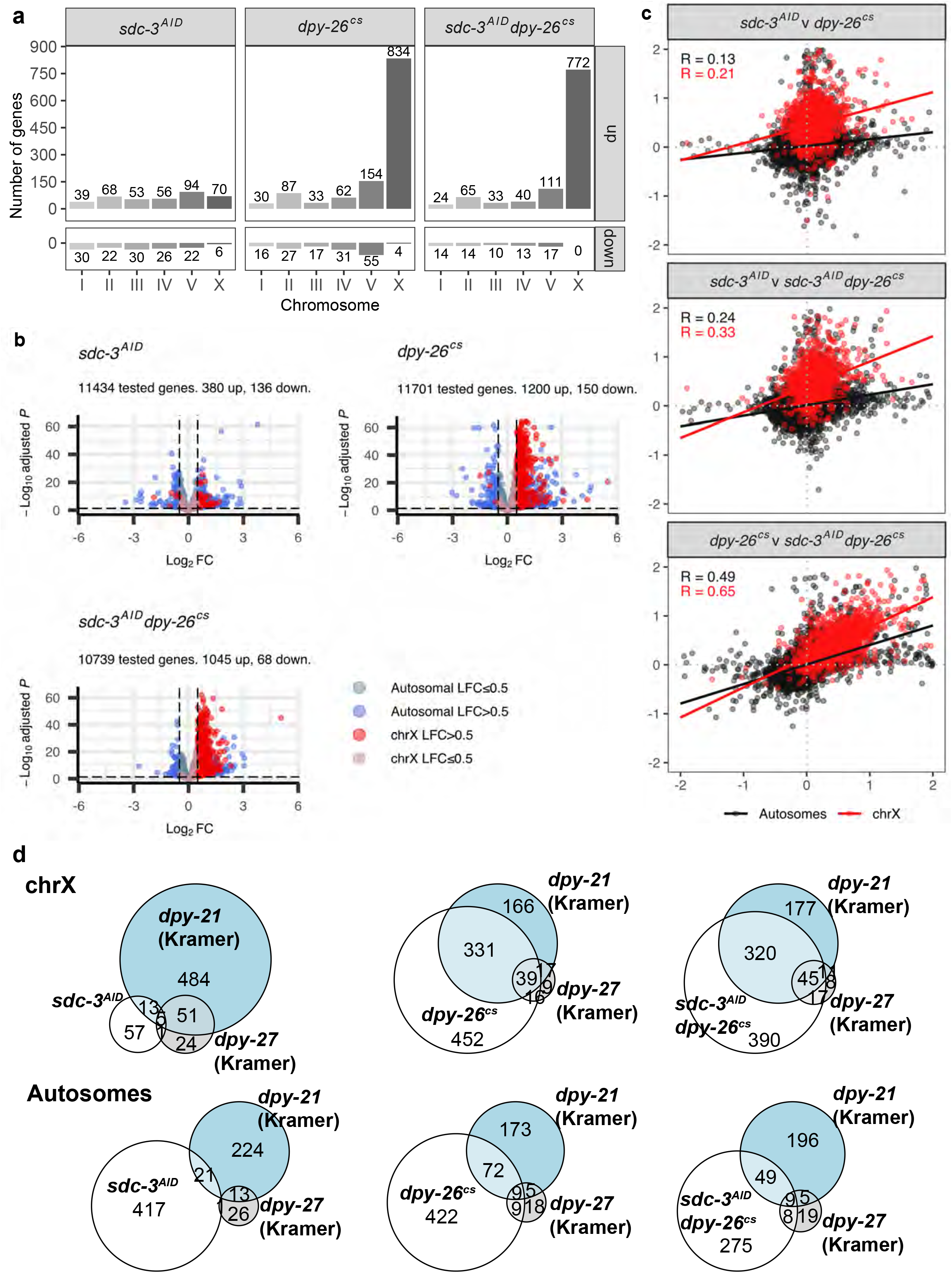
Differential gene expression after inactivation of condensin I/I^DC^ and the SDC complex. **a.** The number of significantly up and down-regulated genes (AdjP<0.05, |LFC|>0.5) after auxin dependant degradation of SDC-3 (subunit of SDC complex required for recruitment of condensin I/I^DC^, *sdc-3^AID^*), cleavage of condensin I/I^DC^ (*dpy-26^cs^*) or combined degradation of SDC-3 (*sdc-3^AID^*) and cleavage of condensin I/I^DC^ (*dpy-26^cs^*). **b.** Volcano plots for all genes filtered by DESeq2 as expressed. X-linked genes are shown in red (|LFC|>0.5) and dull red (|LFC|≤0.5) and autosomal genes are shown in blue (|LFC|>0.5) and dull blue (|LFC|≤0.5). The horizontal dashed line marks an AdjP value threshold of 0.05, and the vertical dashed lines mark LFC thresholds of −0.5 and 0.5. The number of statistically significant (AdjP<0.05, |LFC|>0.5) genes in each dataset is shown in the plot sub-heading. **c.** Pairwise correlation of the log2 fold change of 13734 genes (that have at least 10 read counts in total) upon degradation of SDC-3 (*sdc-3^AID^*), condensin I/I^DC^ (*dpy-26^cs^*) or both (*sdc-3^AID^dpy-26^cs^*). X-linked genes are shown in red, and autosomal genes in black. The lines represent the estimated linear regression, and the Pearson correlation coefficients for autosomes and chrX are shown in each panel. **d.** Overlap between the significantly changing genes in our datasets and previously published gene expression analysis in L3 worms of dosage compensation using RNAi for *dpy-27* (condensin I^DC^) and a *dpy-21(e428)* mutant (SDC complex)^75^. Data from Kramer *et al.* was filtered to remove oscillating genes, and significance was determined at the same AdjP and LFC thresholds as our data (AdjP<0.05, |LFC|>0.5). Data for *dpy-26^cs^* is the same as in Fig. 5, included here for comparison.

## Supplementary tables

**Supplementary Table S1.**
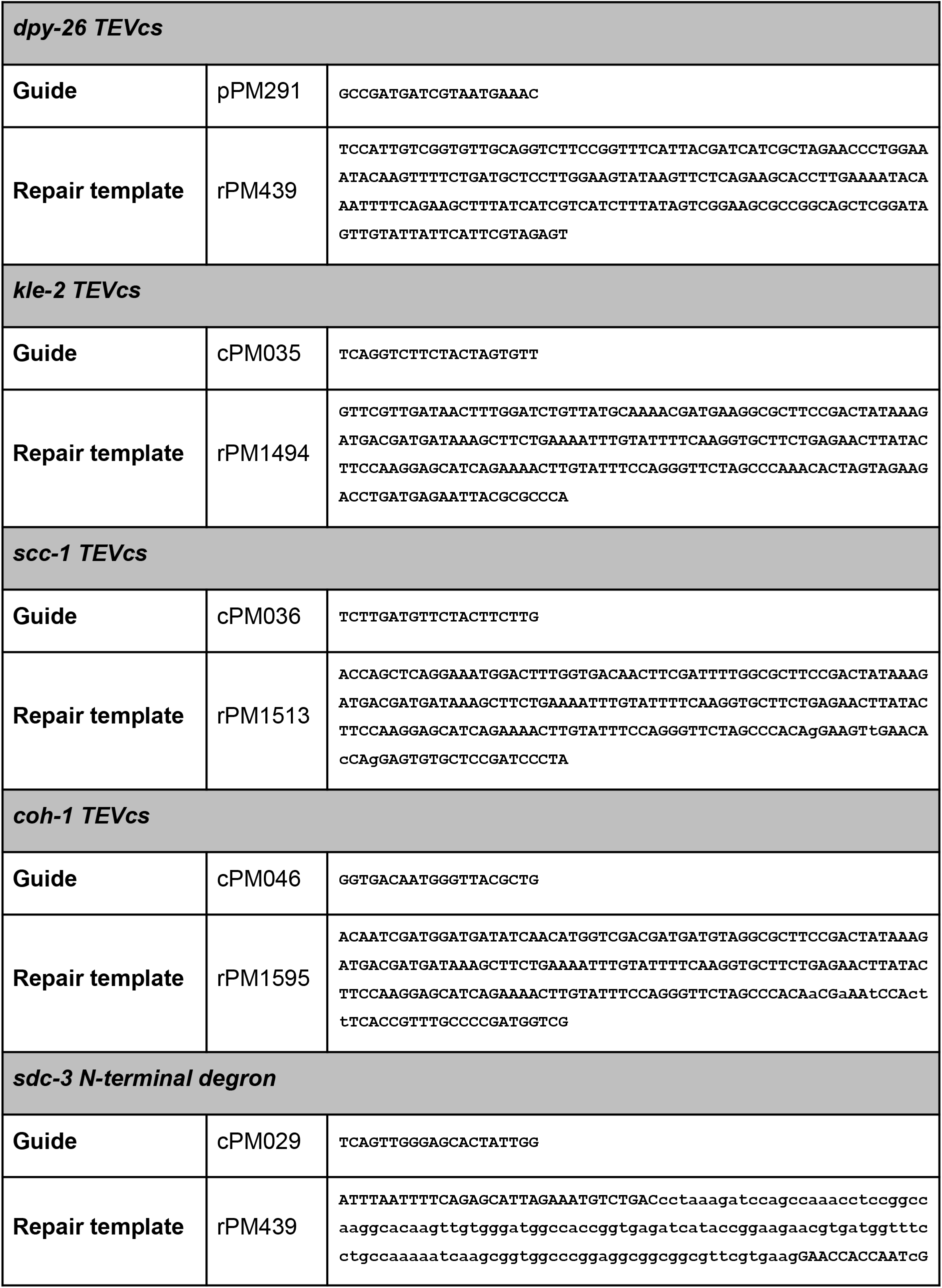
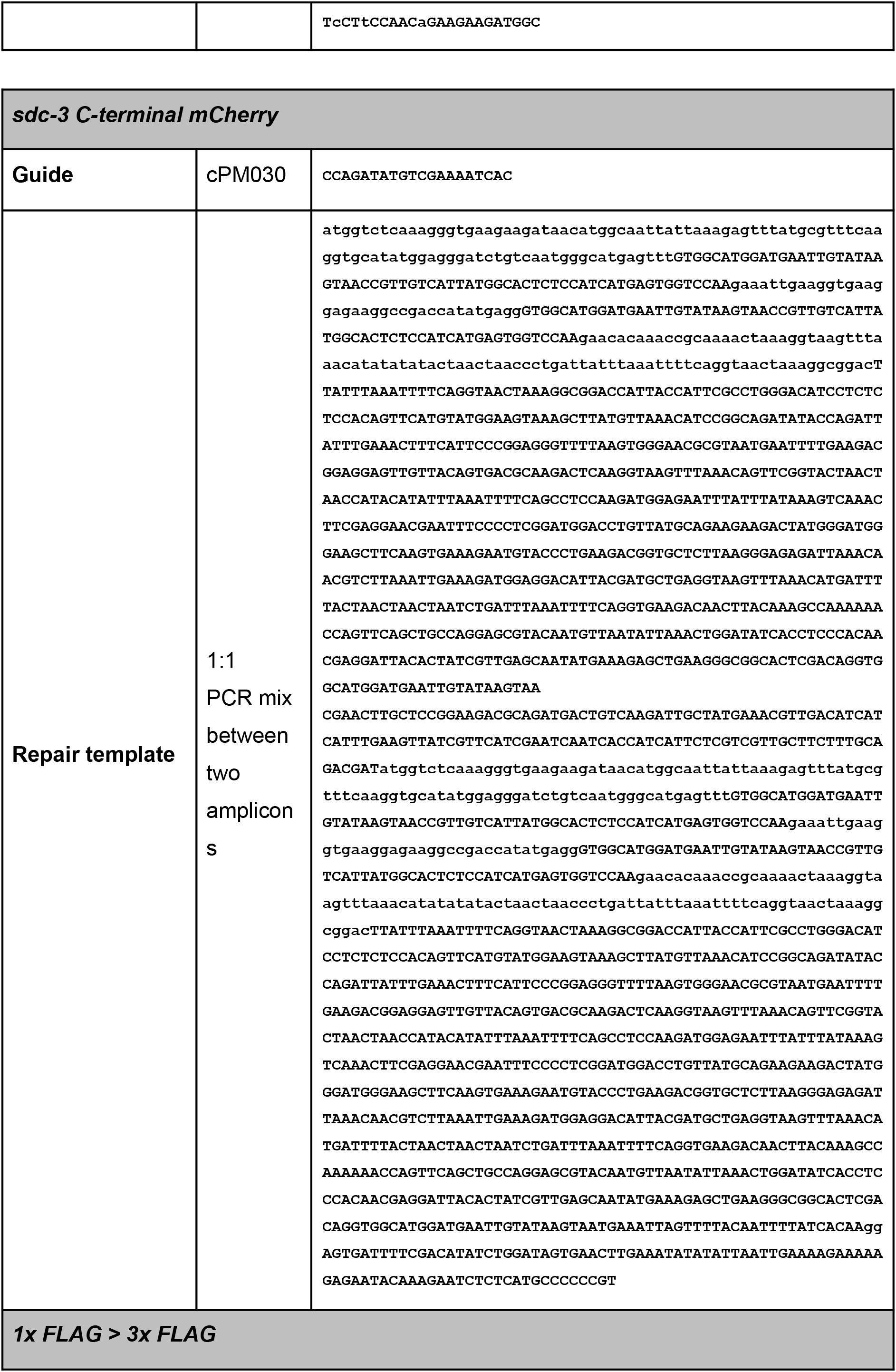
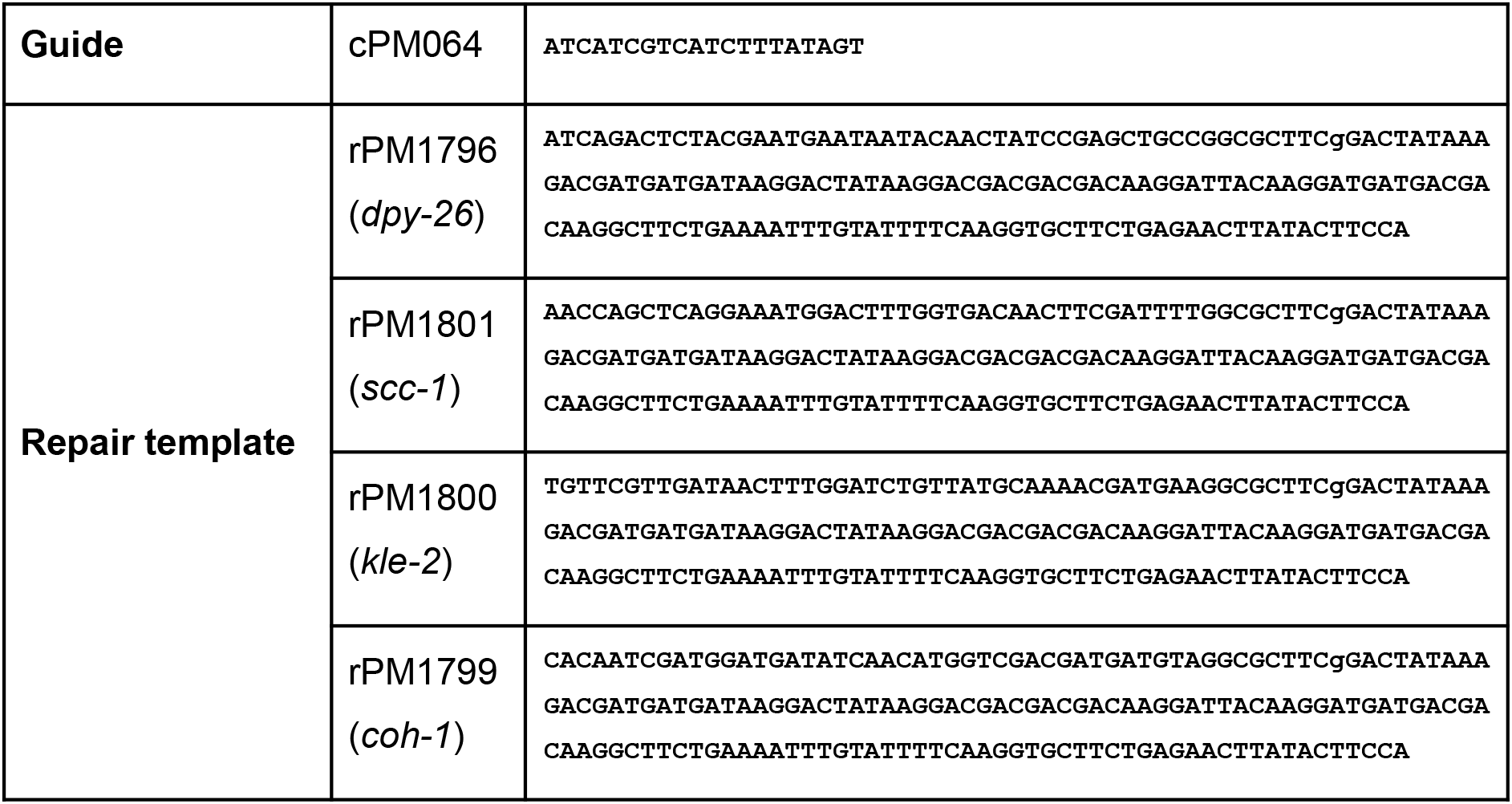
guides and repair templates for kleisin TEV cut-site integration and degron tagging.

**Supplementary Table S2.** *C. elegans* strains and genotypes.

| **Worm strain** | **Genotype** |
| --- | --- |
| PMW366 | *ubsSi20[hsp-16.2p::TEV] II; unc-119(?) III* |
| PMW382 | *ubsSi20[hsp-16.2p::TEV] II; unc-119(?) III; dpy-26(ubs43[1xFLAG TEVcs]) IV* |
| PMW775 | *ubsSi20[hsp-16.2p::TEV] II; unc-119(?) kle-2(ubs14[1xFLAG TEVcs]) III;* |
| PMW784 | *scc-1(ubs19[1xFLAG TEVcs]) ubsSi20[hsp-16.2p::TEV] II; unc-119(?) III* |
| PMW788 | *xeSi376[Peft-3::TIR1::mRuby] unc-119(?) III; sdc-3(ubs20[N-term degraon]) V* |
| PMW821 | *ubsSi20[hsp-16.2p::TEV] II; xeSi376[eft-3p::TIR1::mRuby] III; dpy-26(ubs43[1xFLAG TEVcs]) IV; sdc-3(ubs20[N-term AID]) V.* |
| PMW822 | *ubsSi20[hsp-16.2p::TEV] II; xeSi376[eft-3p::TIR1::mRuby]; III; sdc-3(ubs20[N-term degraon]) V.* |
| PMW823 | *ubsSi20[hsp-16.2p::TEV] II; xeSi376[eft-3p::TIR1::mRuby] III.* |
| PMW828 | *ubsSi20[hsp-16.2p::TEV] II; unc-119(?) III; coh-1(ubs26[1xFLAG TEVcs]) X* |

| Worm strain | Genotype |
| --- | --- |
| PMW831 | *ubsSi20[hsp-16.2p::TEV] II; unc-119(?) III; wls54[scm::gfp] V;* |
| PMW832 | *ubsSi20[hsp-16.2p::TEV] II; unc-119(?) III; dpy-26(ubs43[1xFLAG TEVcs]) IV; wls54[scm::gfp] V* |
| PMW834 | *ubsSi20[hsp-16.2p::TEV] II; unc-119(?) kle-2(ubs14[1xFLAG TEVcs]) III; wls54[scm::gfp] V* |
| PMW836 | *scc-1(ubs19[1xFLAG TEVcs]) ubsSi20[hsp-16.2p::TEV] II; unc-119(?) III; wls54[scm::gfp] V* |
| PMW844 | *scc-1(ubs19[1xFLAG TEVcs]) ubsSi20[hsp-16.2p::TEV] II; unc-119(?) III; coh-1(ubs26[1xFLAG TEVcs]) X* |
| PMW853 | *ubsSi20[hsp-16.2p::TEV] II; unc-119(?) III; wls54[scm::gfp] V; coh-1(ubs26[1xFLAG TEVcs]) X* |
| PMW864 | *scc-1(ubs19[1xFLAG TEVcs]) ubsSi20[hsp-16.2p::TEV] II; unc-119(?) III; wls54[scm::gfp] V; coh-1(ubs26[1xFLAG TEVcs]) X* |
| PMW929 | *wrdSi23[eft-3p::TIR1::F2A::BFP::AID\*::NLS::tbb-2 3'UTR] I; sdc-3(ubs20[N-term degron] ubs39[C-term mCherry])V; yglIs[Imn-1::GFP::Imn-1 unc-119(+)] X* |
| PMW989 | *bqSi142[pBN20(unc-119(+)) Pemr-1::emr-1::mCherry] II sdc-1(st12225[sdc-1::TY1::EGFP::3xFLAG])X.* |
| PMW1005 | *ubsSi20[hsp-16.2p::TEV::unc-54 3'UTR; Cb-unc-119] II; unc-119(?) III; dpy-26[ubs43 ubs44 3xFLAG TEVcs] IV.* |
| PMW1021 | *scc-1(ubs19 ubs46 [3xFLAG TEVcs]), ubsSi20[hsp-16.2p::TEV::unc-54 3'UTR; Cb-unc-119] II; unc-119(?) III.* |
| PMW1023 | *ubsSi20[hsp-16.2p::TEV::unc-54 3'UTR; Cb-unc-119] II; unc-119(?) kle-2(ubs14 ubs48 [3x FLAG TEVcs]) III.* |
| PMW1025 | *ubsSi20[hsp-16.2p::TEV::unc-54 3'UTR; Cb-unc-119] II; coh-1(ubs26 ubs50 [3x FLAG TEVcs]) X.* |

**Supplementary Table S3.** HiC datasets.

| **Strain** | **Initial paired reads** | **Final HiC contacts** | **Trans contacts** | **Cis contacts** |
| --- | --- | --- | --- | --- |
| N2 L3 animals<br>WT_replicate1 | 651'568'455 | 206'920'752 | 27'008'011 | 179'912'741 |
| N2 L3 animals<br>WT_replicate2 | 221'461'300 | 139'711'527 | 18'188'942 | 121'522'585 |
| TEV control,<br>L3 animals, HS<br>PMW366_replicate1 | 459'792'268 | 258'355'543 | 26'902'516 | 231'453'027 |
| TEV control,<br>L3 animals, HS<br>PMW366_replicate2 | 257'937'617 | 163'424'616 | 15'141'862 | 148'282'754 |
| *dpy-26<sup>cs</sup>* TEV,<br>L3 animals, HS<br>PMW382_replicate1 | 361'675'062 | 223'887'805 | 44'083'298 | 179'804'507 |
| *dpy-26<sup>cs</sup>* TEV,<br>L3 animals, HS<br>PMW382_replicate2 | 230'552'312 | 139'901'094 | 29'977'318 | 109'923'776 |
| *kle-2<sup>cs</sup>* TEV,<br>L3 animals, HS<br>PMW775_replicate1 | 318'976'878 | 204'931'744 | 19'515'846 | 185'415'898 |
| *kle-2<sup>cs</sup>* TEV,<br>L3 animals, HS<br>PMW775_replicate2 | 232'861'204 | 154'903'072 | 14'979'338 | 139'923'734 |
| *scc-1<sup>cs</sup>* TEV,<br>L3 animals, HS<br>PMW784_replicate1 | 507'758'707 | 317'973'695 | 41'022'043 | 276'951'652 |
| *scc-1<sup>cs</sup>* TEV,<br>L3 animals, HS<br>PMW784_replicate2 | 415'575'243 | 250'444'362 | 25'432'942 | 225'011'420 |
| *coh-1<sup>cs</sup>* TEV, | 370'162'970 | 208'298'545 | 22'789'025 | 185'509'520 |

| Strain | Initial paired reads | Final HiC contacts | Trans contacts | Cis contacts |
| --- | --- | --- | --- | --- |
| L3 animals, HS<br>PMW828_replicate1 |  |  |  |  |
| *coh-1<sup>CS</sup>* TEV,<br>L3 animals, HS<br>PMW828_replicate2 | 207'132'085 | 130'155'406 | 13'841'786 | 116'313'620 |
| *scc-1<sup>CS</sup> coh-1<sup>CS</sup>* TEV,<br>L3 animals, HS<br>PMW844_replicate1 | 287'910'802 | 172'186'520 | 18'968'929 | 153'217'591 |
| *scc-1<sup>CS</sup> coh-1<sup>CS</sup>* TEV,<br>L3 animals, HS<br>PMW844_replicate2 | 237'756'765 | 148'507'656 | 15'461'079 | 133'046'577 |
| *sdc-3<sup>AID</sup>* N-terminal degron<br>L3 animals, HS<br>PMW822_replicate1 | 344'081'627 | 188'922'870 | 21'746'968 | 167'175'902 |
| *sdc-3<sup>AID</sup>* N-terminal degron<br>L3 animals, HS<br>PMW822_replicate2 | 216'499'112 | 134'778'142 | 17'199'892 | 117'578'250 |

**Supplementary Table S4.** number of animals per lifespan.

|  | Lifespan 1 | Lifespan 2 | Lifespan 3 |
| --- | --- | --- | --- |
| Strain/condition | n assayed<br>(n total) | n assayed<br>(n total) | n assayed<br>(n total) |
| TEV control, PMW366 (HS) | 220 (229) | 332 (436) | 198 (248) |
| *dpy-26* TEV, PMW382 (HS) | 417 (421) | 132 (156) | 61 (70) |
| *kle-2* TEV, PMW775 (HS) | 304 (316) | 248 (256) | 235 (241) |
| *scc-1* TEV, PMW784 (HS) | 254 (262) | 361 (367) | 283 (289) |
| *sdc-3* N-terminal degron,<br>PMW788 (1mM auxin) | 457 (463) | 484 (496) | 461 (475) |
| TEV control males healthspan,<br>PMW366 (HS) | 200 (200) | 175 (200) | 150 (200) |
| *dpy-26* TEV males healthspan,<br>PMW382 (HS) | 200 (200) | 175 (200) | 150 (200) |

**Supplementary Table S5.**
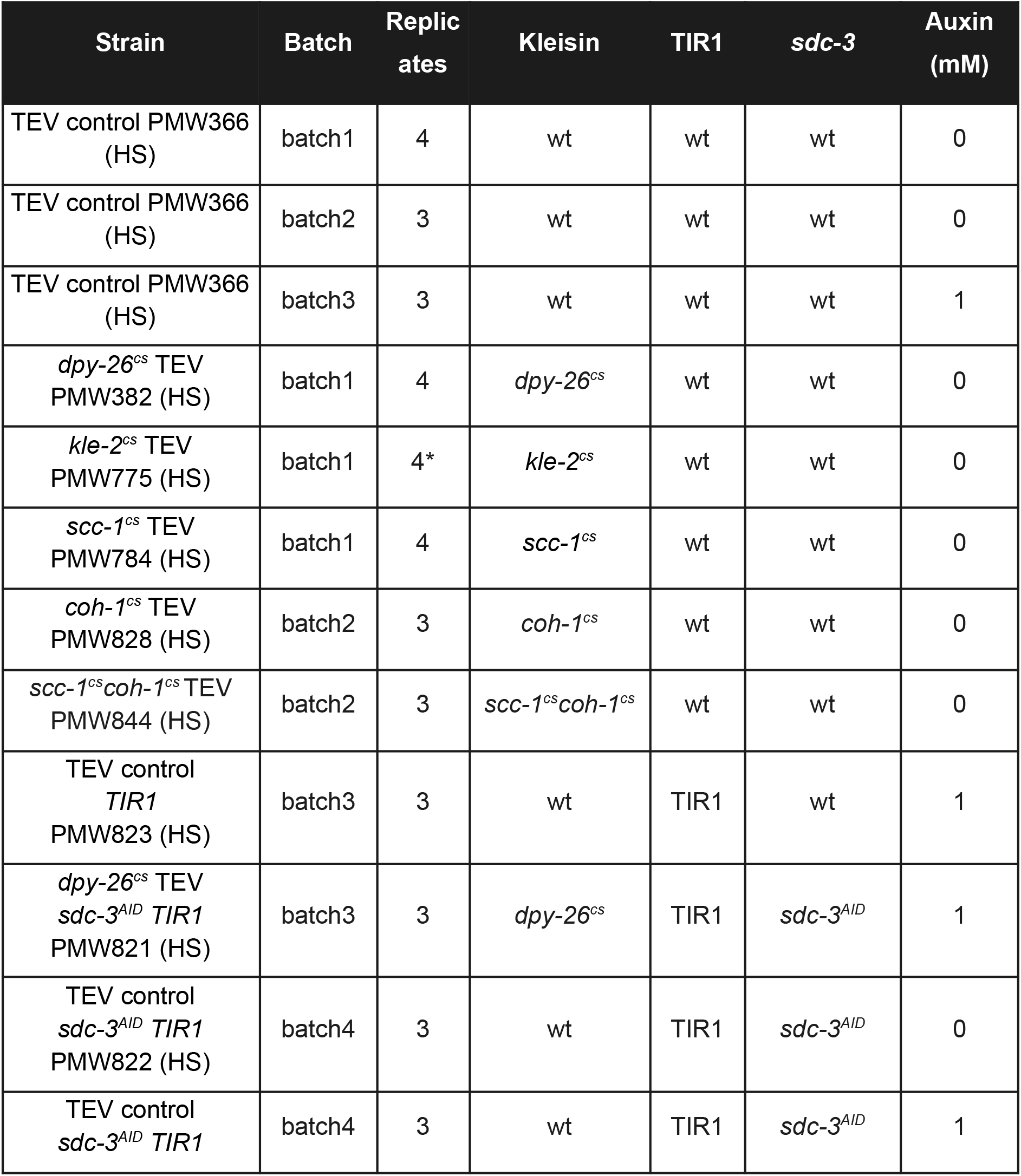
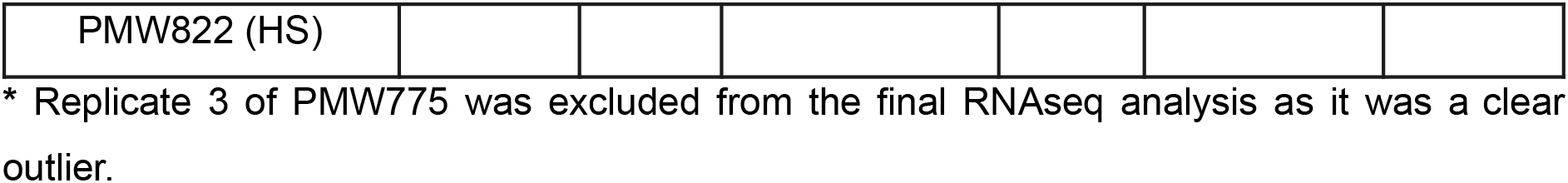
RNAseq samples.

RNAseq samples
| Strain | Batch | Replicates | Kleisin | TIR1 | *sdc-3* | Auxin (mM) |
| --- | --- | --- | --- | --- | --- | --- |
| TEV control PMW366 (HS) | batch1 | 4 | wt | wt | wt | 0 |
| TEV control PMW366 (HS) | batch2 | 3 | wt | wt | wt | 0 |
| TEV control PMW366 (HS) | batch3 | 3 | wt | wt | wt | 1 |
| *dpy-26<sup>cs</sup>* TEV PMW382 (HS) | batch1 | 4 | *dpy-26<sup>cs</sup>* | wt | wt | 0 |
| *kle-2<sup>cs</sup>* TEV PMW775 (HS) | batch1 | 4* | *kle-2<sup>cs</sup>* | wt | wt | 0 |
| *scc-1<sup>cs</sup>* TEV PMW784 (HS) | batch1 | 4 | *scc-1<sup>cs</sup>* | wt | wt | 0 |
| *coh-1<sup>cs</sup>* TEV PMW828 (HS) | batch2 | 3 | *coh-1<sup>cs</sup>* | wt | wt | 0 |
| *scc-1<sup>cs</sup> coh-1<sup>cs</sup>* TEV PMW844 (HS) | batch2 | 3 | *scc-1<sup>cs</sup> coh-1<sup>cs</sup>* | wt | wt | 0 |
| TEV control *TIR1* PMW823 (HS) | batch3 | 3 | wt | TIR1 | wt | 1 |
| *dpy-26<sup>cs</sup>* TEV *sdc-3<sup>AID</sup> TIR1* PMW821 (HS) | batch3 | 3 | *dpy-26<sup>cs</sup>* | TIR1 | *sdc-3<sup>AID</sup>* | 1 |
| TEV control *sdc-3<sup>AID</sup> TIR1* PMW822 (HS) | batch4 | 3 | wt | TIR1 | *sdc-3<sup>AID</sup>* | 0 |
| TEV control *sdc-3<sup>AID</sup> TIR1* | batch4 | 3 | wt | TIR1 | *sdc-3<sup>AID</sup>* | 1 |
| PMW822 (HS) |  |  |  |  |  |  |

\* Replicate 3 of PMW775 was excluded from the final RNAseq analysis as it was a clear outlier.

**Supplementary Table S6.** loop anchors.

| Loop anchors bins |  |  |  |  | rex sites (Albritton et al. 2017 PMID 28562241) |  |  |  |
| --- | --- | --- | --- | --- | --- | --- | --- | --- |
| chr | start | end | name | strength | chr | start | end | name |
| chrX | 410000 | 412000 | anchor_1 | 2.633844179 | chrX | 409986 | 410388 | rex-26_Prex-1 |
| chrX | 490000 | 492000 | anchor_2 | 0.970991956 | chrX | 489659 | 490061 | rex-62 |
| chrX | 784000 | 786000 | anchor_3 | 2.284551637 | chrX | 785349 | 785751 | rex-46 |
| chrX | 806000 | 808000 | anchor_4 | 2.596722342 | chrX | 806476 | 806878 | rex-5_rex-40 |
| chrX | 1346000 | 1348000 | anchor_5 | 2.218487611 | chrX | 1344808 | 1345210 | rex-16_rex-45 |
| chrX | 1378000 | 1380000 | anchor_6 | 1.69569072 | chrX | 1379475 | 1379877 | rex-37_rex-18 |
| chrX | 1436000 | 1438000 | anchor_7 | 1.066193335 | chrX | 1437088 | 1437490 | rex-38 |
|  |  |  |  |  | chrX | 1438905 | 1439307 | rex-59 |
| chrX | 1626000 | 1628000 | anchor_8 | 1.917478463 | chrX | 1626919 | 1627321 | rex-14_Prex-7 |
| chrX | 1908000 | 1910000 | anchor_10 | 2.158962952 | chrX | 1908792 | 1909194 | rex-45 |
| chrX | 2364000 | 2366000 | anchor_11 | 1.157541489 | chrX | 2364709 | 2365111 | rex-36 |
| chrX | 2996000 | 2998000 | anchor_12 | 3.159018191 | chrX | 2996899 | 2997301 | rex-6_rex-32 |
| chrX | 4208000 | 4210000 | anchor_13 | 2.653750907 | chrX | 4208539 | 4208941 | rex-40 |
|  |  |  |  |  | chrX | 4209083 | 4209485 | rex-2_rex-23 |
| chrX | 4394000 | 4396000 | anchor_14 | 1.946209636 | chrX | 4395443 | 4395845 | rex-23_rex-1 |
| chrX | 5428000 | 5430000 | anchor_15 | 3.041041959 | chrX | 5429336 | 5429738 | rex-7_rex-34 |
| chrX | 6296000 | 6298000 | anchor_16 | 3.258246712 | chrX | 6296350 | 6296752 | rex-4_rex-33 |
| chrX | 7180000 | 7182000 | anchor_17 | 1.427207488 | chrX | 7181607 | 7182009 | rex-19_rex-24 |
| chrX | 8036000 | 8038000 | anchor_18 | 2.81452002 | chrX | 8036226 | 8036628 | rex-11_rex-14 |
|  |  |  |  |  | chrX | 8048643 | 8049045 | rex-57_rex-17 |
| chrX | 9464000 | 9466000 | anchor_19 | 2.741367593 | chrX | 9465660 | 9466062 | rex-13_rex-47 |
|  |  |  |  |  | chrX | 9467607 | 9468009 | rex-47 |
| chrX | 11094000 | 11096000 | anchor_20 | 3.225511797 | chrX | 11094000 | 11094402 | rex-1_rex-8 |
|  |  |  |  |  | chrX | 11095445 | 11095847 | rex-39 |
| chrX | 11522000 | 11524000 | anchor_21 | 1.18731798 |  |  |  |  |
| chrX | 11898000 | 11900000 | anchor_22 | 1.640278186 | chrX | 11899062 | 11899464 | rex-33_rex-36 |
| chrX | 11936000 | 11938000 | anchor_23 | 2.837534281 | chrX | 11937330 | 11937732 | rex-9_rex-16 |
|  |  |  |  |  | chrX | 11934934 | 11935336 | rex-50 |
| chrX | 12362000 | 12364000 | anchor_24 | 1.927906752 | chrX | 12362973 | 12363375 | rex-17_rex-6 |
| chrX | 13442000 | 13444000 | anchor_25 | 1.874477715 | chrX | 13442434 | 13442836 | rex-41 |
|  |  |  |  |  | chrX | 13435108 | 13435510 | rex-48 |
| chrX | 13512000 | 13514000 | anchor_26 | 2.016204254 | chrX | 13514381 | 13514783 | rex-22_Prex-22 |
| chrX | 13700000 | 13702000 | anchor_27 | 2.513424824 | chrX | 13700665 | 13701067 | rex-3_rex-43 |
|  |  |  |  |  | chrX | 13665390 | 13665792 | rex-52 |
| chrX | 14526000 | 14528000 | anchor_28 | 2.947047809 | chrX | 14525682 | 14526084 | rex-8_Prex-28 |
| chrX | 14814000 | 14816000 | anchor_29 | 1.459982746 | chrX | 14813358 | 14813760 | rex-28_rex-39 |
| chrX | 15736000 | 15738000 | anchor_30 | 1.408724818 | chrX | 15736449 | 15736851 | rex-30_rex-46 |
|  |  |  |  |  | chrX | 15741843 | 15742245 | rex-49 |
| chrX | 16056000 | 16058000 | anchor_31 | 2.634334966 | chrX | 16056410 | 16056812 | rex-15_Prex-30 |
| chrX | 16208000 | 16210000 | anchor_32 | 2.572346009 | chrX | 16209687 | 16210089 | rex-31 |
|  |  |  |  |  | chrX | 16205802 | 16206204 | rex-24_Prex-31 |
|  |  |  |  |  | chrX | 16202289 | 16202691 | rex-44 |
| chrX | 16328000 | 16330000 | anchor_34 | 0.891481062 | chrX | 16327264 | 16327666 | rex-43 |
| chrX | 16682000 | 16684000 | anchor_35 | 2.78131332 | chrX | 16681893 | 16682295 | rex-10_rex-35 |
|  |  |  |  |  | chrX | 16695171 | 16695573 | rex-63 |
| chrX | 17180000 | 17182000 | anchor_36 | 2.877223158 | chrX | 17181251 | 17181653 | rex-21_rex-42 |
| chrX | 17546000 | 17548000 | anchor_37 | 3.440073192 | chrX | 17544344 | 17544746 | rex-12_rex-41 |
|  |  |  |  |  | chrX | 17545518 | 17545920 | rex-20 |

**Note:** Manual inspection of the second *cis* eigenvector of chromosome X upon condensin I/II<sup>DC</sup> cleavage revealed that anchors 9 and 33 are artefacts (local maxima) due to low mapping regions. For consistency, the peak numbering has been conserved, but anchors 9 and 33 were removed.

**Supplementary File 1 - DESeq2 results for *dpy-26^cs^*, *kle-2^cs^*, *scc-1^cs^* and *coh-1^cs^* cleavage and SDC-3^AID^ degradation.** Tables of RNAseq differential expression analysis by DESeq2 for all genes with at least 10 read counts in total among all samples. DESeq2 performs additional filtering based on the baseMean expression values with the threshold determined empirically from the data, resulting in the adjusted p-value of the genes that do not pass this threshold being assigned NA. The adjusted p-values that were below the 0.05 threshold are highlighted in red. Genes with LFC>0.5 are highlighted in green, and genes with a LFC< −0.5 are highlighted in yellow.

**Supplementary File 2 - GO term, tissue and phenotype enrichment analysis for up and down regulated genes upon SMC complex inactivation.** Tables of results of gene ontology (GO), tissue and phenotype enrichment analysis performed with the command line version of the Wormbase EA tool ^115^. Lists of significantly up and down (adjusted p-value < 0.05 and LFC>0.5 or LFC< −0.5) were analyzed separately, but the results were combined in a single table per dataset, with the last column (“UpOrDownRegulated”) indicating the source of the results. The “category” column and the color of the cells indicate whether the enrichment analysis was carried out for GO terms (red), tissue (yellow), or phenotype (green).

